# Development of three-color FRET measurement of force-dependent sensing of RIAM-Vinculin interactions in focal adhesions

**DOI:** 10.1101/2025.03.04.641466

**Authors:** Conor A. Treacy, Tommy L. Pallett, Tam Bui, Simon P. Poland, Mark A. Pfuhl, Maddy Parsons, Simon M. Ameer-Beg

**Affiliations:** Randall Centre for Cell and Molecular Biophysics, King’s College London, UK; Comprehensive Cancer Centre, School of Cancer and Pharmaceutical Sciences, King’s College London, UK; Institute of Pharmaceutical Science, School of Cancer and Pharmaceutical Sciences, King’s College London, UK

**Author notes:** Corresponding Author Simon M. Ameer-Beg.

**Keywords:** Three-color FRET, TCSPC-FLIM, Vinculin, RIAM, Focal Adhesions

## Abstract

Förster resonance energy transfer (FRET) is a powerful technique for probing molecular interactions and conformational changes in biological systems. Cascade-FRET, a multistep energy transfer system involving three fluorophores, enables spatial and temporal mapping of molecular interactions. Here, we leveraged Cascade-FRET with time-correlated single photon counting fluorescence lifetime imaging microscopy (TCSPC-FLIM) to explore the putative interaction between Rap1-interacting Adapter Molecule (RIAM) and vinculin in focal adhesions. We developed a novel three-fluorophore Cascade-FRET system connected by flexible peptide linkers comprising mTurquoise2, mVenus, and mScarlet-I, validated using purified proteins, spectroscopic analysis, structural modeling, and negative staining transmission electron microscopy (TEM). Putative RIAM-vinculin interactions were explored in vinculin knockout mouse embryonic fibroblasts and revealed that RIAM binds to the N-terminus of vinculin in focal adhesions. This complex requires an intact microtubule cytoskeleton. Vinculin tension-sensing constructs quantified intracellular forces, with an average force of 2.95 ± 0.97 pN per focal adhesion. These findings corroborate the mechanosensitive role of vinculin and its interaction with RIAM in a force-independent manner. This study demonstrates the utility of Cascade-FRET and TCSPC-FLIM in investigating multicomponent molecular complexes. Our findings provide novel insights into RIAM-vinculin interactions and their regulation by intracellular tension, paving the way for advanced applications of Cascade-FRET in dynamic cellular systems.

**Significance Statement:** Focal adhesions are used by all adherent cells to attach to their surroundings and transmit mechanical signals. This paper uses a biosensor to measure force changes within the focal adhesion-associated protein called vinculin. The fluorescence lifetime of the biosensor changes when force is applied across vinculin, allowing us to report on changes in force within developing focal adhesions. We show that another protein, RIAM, interacts with vinculin in a force-independent manner. This was achieved by developing a three-color cascade FRET model, showing how the three proteins—talin, vinculin, and RIAM—interact over time. This research has furthered our understanding of the order and mechanism in which these components assemble in cell adhesions.

## Introduction

Förster resonance energy transfer (FRET) techniques applied in cell biology provide data on macromolecular function, structure, dynamics, and the local fluorophore environment^1–3^. FRET can be used as a quantitative proximity sensor between fluorescently labeled proteins with high sensitivity between 1-9 nm^3,4^. This enables the detection of direct intra- and intermolecular interactions by various modalities in fixed and living cells^5^. We employed fluorescence lifetime imaging microscopy (FLIM) using Time-Correlated Single Photon Counting (TCSPC), which provides accurate and sensitive FRET measurements^6–8^. Recent advancements, including faster acquisition speeds, have enabled real-time imaging of dynamic biological processes^9–13^ This has facilitated the development of more sophisticated FRET systems, incorporating multiplexed and cascaded donor-acceptor pairs, which have been investigated at the single-molecule level using ratiometric measurements^14–16^. Three-fluorophore systems comprising two donor-acceptor pairs wherein the first acceptor is the donor in the second FRET pair have been demonstrated^15,17^ (Figure 1A). We are the first to describe this type of FRET model as Cascade-FRET, but similar variants have been used previously for ensemble measurements^17–19^ and, more recently, in single-molecule applications^15,20,21^. The sequential action of cascading multiple FRET interactions has been previously used to investigate multiple conformational states of a single protein^5,22^. Cascade-FRET is not just limited to determining whether an interaction has occurred^14,15^. Recent studies have shown that attachment of three fluorescent dyes to a single protein^14,21,23,24^ or nucleic acid^1,15,25^ enables the determination of the three-dimensional conformation, orientation, and activation state during dynamic processes such as protein folding, ligand binding, or post-translational modification. Furthermore, cascade-FRET can uniquely elucidate the order in which multi-component complexes assemble, interact, evolve, and ultimately disassemble.

**Figure 1.**
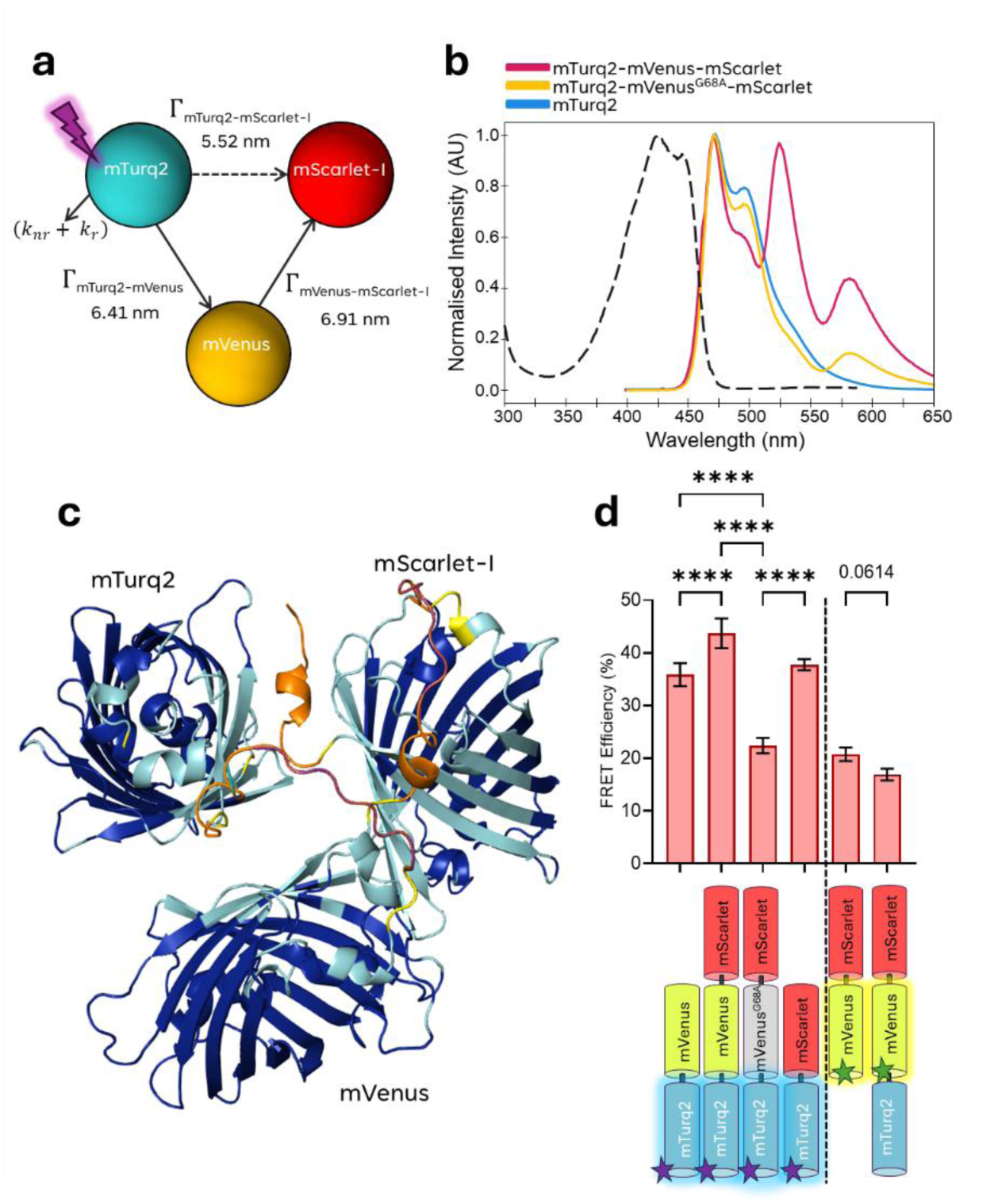
Three-color FRET-Cascade construct. a) A graphical representation of the fluorophores used in the three-color FRET cascade module with specific energy transfers (Γ) labelled. The purple lightning bolt represents excitation at 435 nm. Solid arrows show direction of energy cascade through mVenus, dashed arrow shows mTurq2 to mScarlet-I direct energy transfer. b) Excitation spectra for mTurq2 and the emission spectra for mTurq2 (blue), mTurq2-mVenusG68A-mScarlet-I (yellow), and mTurq2-mVenus-mScarlet-I (red) where mTurq2 alone is excited at 435 nm (excitation spectrum indicated by black dashed line). c) An AlphaFold3 model of the three-color protein complex, regions with a high predicted local distance difference test (pLDDT) score >90 are a dark blue, 70 > 90 light blue, 50 > 70, yellow and those with a very low pLDDT score (< 50) are indicated in orange. d) Average FRET efficiencies for the various FRET pairs measured using our custom TCSPC-FLIM system, n=9 individual cells measured per condition across N=3 technical repeats. Bars are the mean FRET efficiency ± SEM (error bars); comparisons of means are calculated by one-way ANOVA using Dunnet T3 correction for multiple comparisons: P-values ≥ 0.123 ns, ≤ 0.0332 (*), ≤ 0.0021 (**), ≤ 0.0002 (***), ≤ 0.0001 (****), p = 0.0614 (mVenus-mScarlet vs. mTurq2-mVenus-mScarlet).

Focal adhesions (FA) are ideally suited to the application of cascade-FRET. FA are large macromolecular assemblies that anchor cells to the extracellular matrix (ECM) and are the primary site of mechanical force transduction^26,27^. FA consist of integrins, which are heterodimeric transmembrane receptors that directly connect to the ECM via their extracellular ligand-binding domains, as well as numerous intracellular proteins organized into layers which regulate force transduction and interaction with the cytoskeleton^28,29^. Proteins such as talin and kindlin can bind directly to integrin cytoplasmic domains, in part mediated through Rap1-induced membrane-associated protein (RIAM), and mediate recruitment of adaptor and signaling proteins including vinculin, FAK, and paxillin with dynamic linkage to the cytoskeleton ^30–35^. Previous studies have analyzed tension across specific FA proteins to explore the relationship between mechanosensing, FA dynamics and cell migration^36^. The first example of this used a tension sensing module (TSMod) containing a 40-amino acid long elastic domain integrated into vinculin between the Vh and Vt domains, creating a vinculin-tension sensing (VincTS) construct^36,37^. Analysis of the VincTS construct in living cells using FLIM revealed that the average force in stationary focal adhesions was approximately 2.5 pN^36^. A putative binding interaction between the focal adhesion protein vinculin and the adaptor protein RIAM was identified *in vitro* between the N-terminus of RIAM (amino acids 1- 127) and the N-terminus of vinculin (1–258), which was approximately five-fold weaker than the mutually exclusive vinculin-talin interaction^38^. However, the presence of this complex and its function in signaling or mechanosensing in FA remains unclear.

In the present study, we developed a three-color fluorescent protein FRET-Cascade to analyze tripartite protein interactions in live cells, consisting of, mTurquiose2^39^, mVenus^40^, and mScarlet-I^41^. The photophysical properties of this construct were tested *in vitro* and performed predictably; this data was used to reproduce the structure of the model both with purified protein *in vitro* and in cells. Furthermore, introducing a glycine-to-alanine mutation in mVenus at amino acid 68 (Gly68Ala) provided a photochromically dead form of the fluorophore as an optimal control which maintains stoichiometry and structural consistency. We applied this novel FRET-cascade to investigate the activation and binding of the mechano-sensitive protein vinculin with its putative interactor RIAM (Rap-1a Interacting Adaptor Molecule). We characterize the interaction of full-length, fluorescently tagged vinculin and RIAM using transiently transfected vinculin knockout Mouse Embryonic Fibroblasts^36,42^ (MEF^Vinc-/-^) with TCSPC-FLIM. Using a previously published vinculin tension sensor,^37,43^ to explore the mechanical role of vinculin, the previously published vinculin tension sensor^37,43^ (vincTS) was used to measure intracellular forces. We then applied the three-color FRET-cascade model to describe the spatial relationship between vinculin and RIAM in focal adhesions. This work establishes a robust FRET-based approach for studying molecular interactions and intracellular forces. By successfully quantifying RIAM-vinculin binding and vinculin tension, the study provides new insights into focal adhesion mechanics and paves the way for broader applications in mechanobiology.

## Results

### Three-color FRET Cascade Implementation

The generalized theory of stepwise and cascaded FRET efficiency was outlined by *Watrob et al*^16^.; we will describe only the pertinent elements (Figure 1a). The measured donor fluorescence lifetime, 𝜏_𝐷𝐴_, in the presence of an additional FRET decay pathway from the excited state, can be generalized as

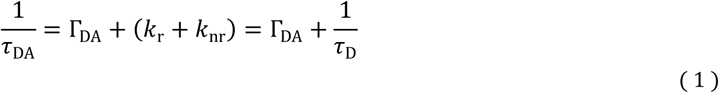

where 𝛤_𝐷𝐴_ is the FRET transfer rate from donor to acceptor, *k* _r_ and *k* _nr_ are the radiative and non-radiative decay rates for the fluorophore, respectively, and 𝜏_D_ the donor lifetime in the absence of FRET. This can be rearranged to give generalized FRET efficiency for donor-acceptor pairs

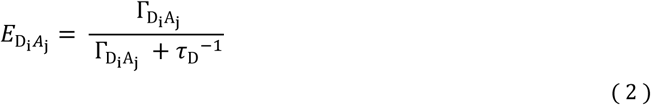

Where the indices *i* and *j* indicate the position of the donor and acceptor in the cascade. Such a system is readily scalable and may contain any number of donors and acceptors. For any such system, total FRET efficiency measured for a given donor, D *_i_* is dependent on a sum over *n* acceptors:

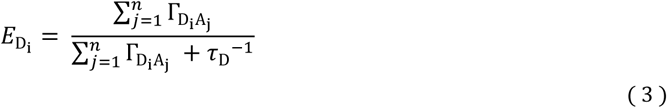

Calculating the individual transfer rates allows us to extract the separation of donor and acceptor pairs directly. Given that the transfer rate is determined as

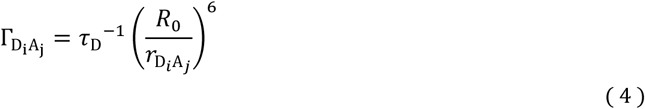

Where R_0_ is the Förster radius and 𝑟_D𝑖A𝑗_ the molecular separation between D *_i_* and A *_j_* . It is possible to dissect the molecular separations not simply on a pairwise basis but, in the presence of multiple interactions, to determine the coordinates of each molecule relative to the donor based on the modified lifetime of D *_i_* alone, given suitable controls. Such a system is generalizable to any number of fluorophores but quickly becomes unwieldy due to error propagation. In this paper, we restrict the experimental verification to 3 fluorophores.

A three-color model to demonstrate the FRET-cascade principle was designed, consisting of mTurquoise2 (mTurq2)^44^, mVenus^40^ and mScarlet-I^41^ (Figure 1a). For each purified protein, excitation and emission spectra are given in Supplementary Figure S1. Each fluorescent protein is separated by a flexible 6 amino acid linker (GGSGGS). Fluorophores act as a donor and/or acceptor within an energy transport cascade. mTurq2 was chosen for its high quantum yield (0.93) and mono-exponential fluorescence emission lifetime of ∼4.1 ns (*in vitro* solution)^39^. mVenus is well documented as an excellent FRET acceptor for mTurq2^39,45^, and mScarlet-I, due to its high absorption cross-section, was chosen as an acceptor to either mTurq2 or mVenus^39,45^. mScarlet-I has substantial spectral overlap with mVenus and mTurq2, with advantageous Forster radius (R_0_) and brightness compared to other red fluorescent proteins^39^. Essential metrics for selecting fluorophores include the Förster radius and the quantum yield of the first acceptor, as these factors ultimately limit stepwise interactions. Optimizing these parameters is particularly advantageous when applying the FRET cascade to living cells, (Supplementary S2 and Table S1).

A full range of pairwise constructs was purified, and the single FP spectra were compared with the mTurq2-mVenus, and mTurq2-mScarlet-I constructs (Supplementary Figure S3). The excitation spectra for mTurq2-mVenus are similar to those for mTurq2 alone because only the mTurq2 is excited at 435 nm. However, the emission spectra differ, particularly at ∼530 nm, corresponding to the peak emission of mVenus. The mTurq2-mVenus construct shows no spectral features corresponding to mScarlet-I as expected. Since mVenus is minimally excited at 435 nm, the emission peak at 530 nm must result from mVenus FRET sensitized emission, with the magnitude of acceptor emission proportional to the FRET efficiency between the two fluorescent proteins (FPs). A similar pattern is observed in the mTurq2-mScarlet-I protein, with the relative heights of these peaks corresponding to the FRET efficiency of the interaction. The emission spectra of mTurq2 alone are compared with those of the three-color protein mTurq2-mVenus-mScarlet-I (Figure 1b). Upon excitation at 435 nm, the emission spectra of the three-color protein display peaks corresponding to all three reference spectra. The prominent peak at 530 nm indicates FRET between mTurq2 and mVenus. An additional peak at approximately 590 nm, corresponding to mScarlet-I, indicates a secondary FRET transition, where energy is transferred from either mTurq2 or mVenus to mScarlet-I, albeit with lower coupling efficiency. The presence of mVenus increases the separation between mTurq2 and mScarlet-I, reducing coupling via FRET compared to the mTurq2-mScarlet-I construct in the absence of mVenus.

A structural control containing a glycine-to-alanine mutation in the fluorochrome of mVenus designated mTurquoise2-mVenus^G68A^-mScarlet-I, was cloned and purified. This mutation prevents mVenus from forming a functional fluorochrome (supplementary methods), therefore serving as an absorption/emission null structural control for the FRET-Cascade model, whilst still properly folding and thereby maintaining the overall structure. Spectral and circular dichroism measurements for mutated mTurquoise2-mVenus^G68A^-mScarlet-I demonstrate the ablated mVenus fluorescence whilst remaining folded and retaining mScarlet-I sensitized emission (Figure 1b, supplementary S4 and supplementary methods).

The three-color FRET-cascade construct was modelled using AlphaFold3^46^ (Figure 1c) using the single full-length sequence of the three-color peptide. AlphaFold3 consistently predicted a triangular 3D tertiary structure formed by the β-barrels of the three FPs, though their relative orientations remain undefined. For further details and model validation, see Supplementary Figures S5, S6 and the supplementary methods.

The three-color proteins, mTurq2-mVenus-mScarlet-I and mTurq2-mVenus^G68A^-mScarlet-I exhibited reductions in fluorescence lifetime compared to their single FP controls (Figure 1d, Supplementary Figure S7). However, mTurq2-mVenus^G68A^-mScarlet-I had a longer fluorescence lifetime, consistent with the lack of a functional mVenus acting as an intermediary in the FRET-cascade. Average lifetimes, FRET efficiencies and R_0_ values are detailed in Table 1. FRET efficiency for the mTurq2-mVenus protein increased with the addition of mScarlet-I indicating FRET between mTurq2 and mScarlet-I.

**Table 1.** Three-color FRET model Lifetimes and FRET efficiencies. Summary tables detailing the fluorescence lifetimes, and χ^2^/degrees of freedom (goodness of fit metric) for the purified proteins used in the FRET cascade model. FRET efficiencies, Forster radii (R_0_), separation distance (R) and their respective SEM are also presented. N=9 measurements cells per condition across three separate technical repeats.

| Purified Protein | Average lifetime (ns) | SEM | Average $\chi^2$/d.f. |
| --- | --- | --- | --- |
| **mTurq2** | 4.178 | 0.01177 | 0.949 |
| **mVenus** | 3.020 | 0.03676 | 1.095 |
| **mTurq2-mVenus** | 2.668 | 0.04371 | 0.908 |
| **mVenus-mScarlet** | 2.393 | 0.02838 | 0.991 |
| **mTurq2-mScarlet** | 2.600 | 0.01738 | 1.089 |
| **mTurq2-mVenus<sup>G68A</sup>-mScarlet** | 3.242 | 0.03438 | 1.159 |
| **mTurq2-mVenus-mScarlet** | 2.332 | 0.05397 | 1.058 |
| **mTurq2-mVenus-mScarlet** | 2.509 | 0.06745 | 1.075 |

| FRET Pair | FRET (%) | SEM | $R_0$ (nm) | R (nm) | SEM |
| --- | --- | --- | --- | --- | --- |
| **mTurq2-mVenus** | 35.89 | 0.564 | 5.83 | 6.41 | 5.23E-02 |
| **mTurq2-mVenus-mScarlet** | 43.72 | 0.892 | n.a. | n.a. | n.a. |
| **mTurq2-mVenus<sup>G68A</sup>-mScarlet** | 22.41 | 0.4572 | 5.08 | 6.25 | 5.49E-02 |
| **mTurq2-mScarlet** | 37.77 | 0.2739 | 5.08 | 5.52 | 2.15E-02 |
| **mVenus-mScarlet** | 20.78 | 0.7336 | 5.53 | 6.91 | 1.03E-01 |
| **mTurq2-mVenus-mScarlet** | 16.94 | 0.6397 | 5.53 | 7.21 | 1.09E-01 |

Energy transfer rates were calculated from fluorescence lifetime data, allowing modeling of a theoretical three-color FRET exchange (Table 2). The calculated FRET transfer rates correspond to distances and average angles between the arms formed by mVenus with mScarlet and mTurq2 that are consistent with the AlphaFold3^47^ model prediction of a triangular configuration (Figure 1c). However, the absolute values of the distances are shorter in the predicted structure from AlphaFold model, likely due to not implementing Amber relaxation to correct for steric hindrances ^48^and because AlphaFold does not account for solvation. Both factors result in a model which is condensed compared to the native setting. These findings, while not conclusive, provide validation for the proposed model (supplementary Figure S8).

**Table 2.** Three-color FRET model energy transfers. a) A summary table detailing the energy transfer rates, SEM, and the associated lifetime of each energy transfer in the construct. K = energy transfer rate for a specific fluorophore; Γ = specific FRET energy transfer within a FRET pair. N=9 measurements cells per condition across three separate technical repeats.

| | Energy Transfer ($s^{-1}$) | SEM | Lifetime (ns) |
| --- | --- | --- | --- |
| $K_{mTuq2}$ | 2.39E+08 | 2.81E-08 | 4.178 |
| $K_{mTuq2-mVenus}$ | 3.75E+08 | 1.64E-07 | 2.668 |
| $\Gamma_{mTuq2-mVenus}$ | 1.35E+08 | 5.91E-08 | 7.382 |
| $K_{mTuq2-mVenusG68A-mScarlet-I}$ | 3.08E+08 | 1.06E-07 | 3.242 |
| $\Gamma_{mTuq2-mScarlet-I}$ | 6.91E+07 | 1.40E-07 | 14.471 |
| $K_{mTuq2-mVenus-mScarlet-I}$ Calculated | 4.44E+08 | 2.32E+07 | 2.253 |
| $K_{mTuq2-mVenus-mScarlet-I}$ Measured | 4.29E+08 | 1.47E+07 | 2.331 |
| Difference (%) |  |  | 3.36% |

Negative Staining Transmission Electron Microscopy (TEM) was used to obtain low-resolution (<1 nm) structural images of the three-color mTurq2-mVenus-mScarlet-I protein (supplementary Figure S9) to validate the FRET-derived intramolecular distances. The TEM images reveal a geometry that matched well with the FRET model, and the distance and angle values agreed within the standard error. This agreement is notable given the simplicity of the FRET model, though some discrepancies may arise from the protein’s orientation on the TEM grid. A comparison of the distances and angles between the calculated FRET, AlphaFold3 model and TEM measurements are detailed in supplementary Table S2.

### Exemplification of the FRET cascade system in a biologically relevant context

Energy transfer rates obtained from the model FRET cascade can be used to determine changes in distances between adjacent biological molecules in a dynamic complex. We employ this method to validate our hypothesis that RIAM binds to vinculin under tension in cells and map the change in applied forces to vinculin on association with RIAM. Whilst the RIAM-vinculin interaction was previously observed *in vitro*^38^, confirmation *in vivo* and a mechanistic description is absent from the literature. Our AlphaFold3 model of the interaction between amino acids 1-30 of RIAM and autoinhibited full-length vinculin (Figure 2a, supplementary Figures S10 and S11) supports published *in vitro* analytical gel filtration data^38^. Our AlphaFold3 model gives further evidence that RIAM and vinculin interact in isolation and when vinculin is in an autoinhibited state.

**Figure 2.**
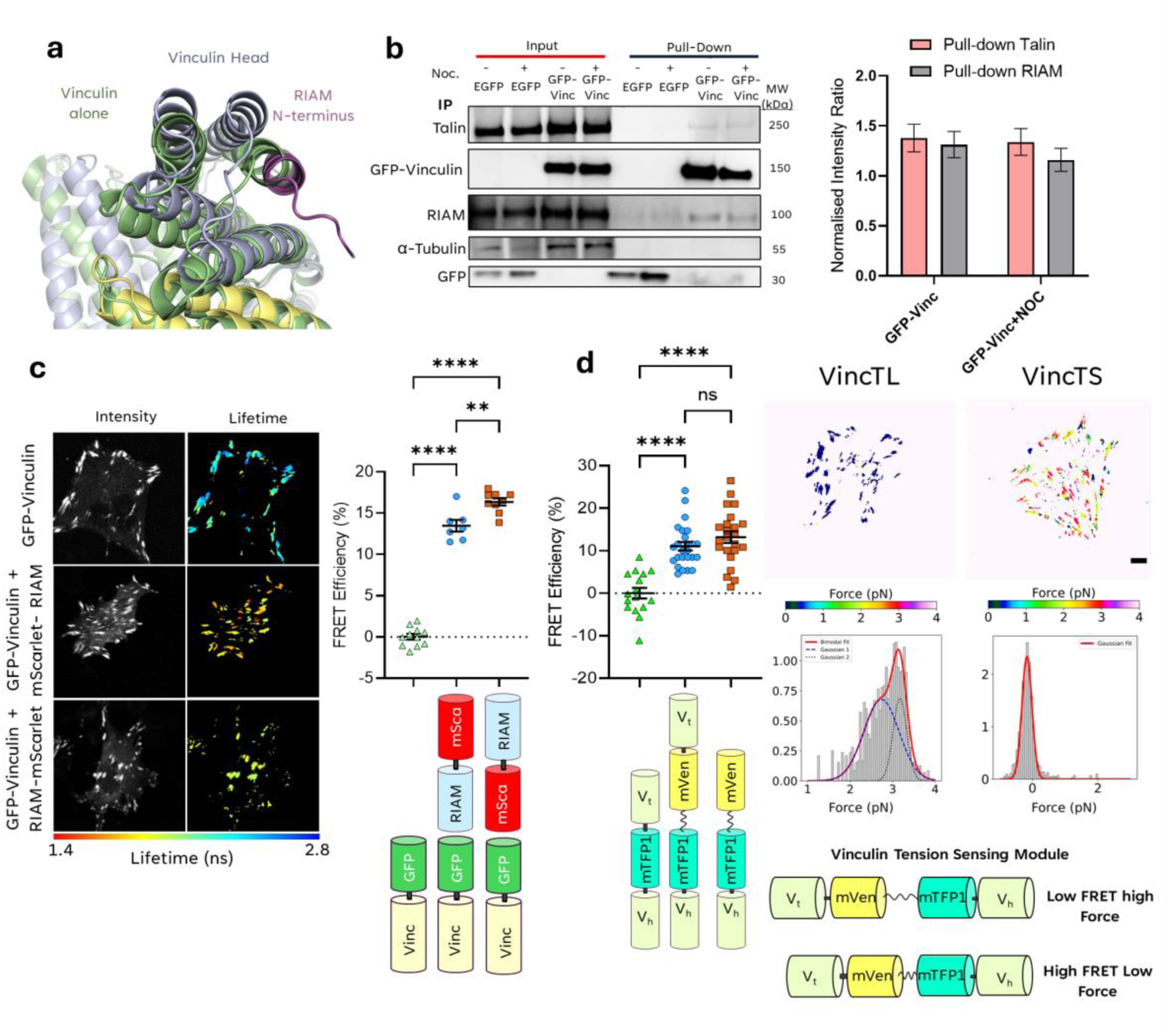
Establishment of the putative RIAM-vinculin interaction. a) An AlphaFold3 model of the RIAM-vinculin interaction. b) A western blot of the co-immunoprecipitation of EGFP-vinculin with RIAM and Talin, probed with an antibody against EGFP. Noc. = Nocodazole treatment. To the right of the blot is a quantification of the pixel intensities of the protein bands for Talin and RIAM pulled down by GFP-Vinculin. The bar graph represents the mean intensity normalized to the total protein with error bars representing the standard error in the mean. Data taken from N=3 blots; the blot displayed is a representative example. c) Fluorescence lifetime data for EGFP-vinculin ± mScarlet-RIAM/RIAM-mScarlet with average FRET efficiencies plot alongside. d) Fluorescence lifetime data for vincTS and controls. The right-hand panel shows typical FRET efficiency distributions and associated tension distributions for a vincTL or vincTS transfected MEF^Vinc-/-^. Above are color-scaled force maps; scale bar = 5 µm. Below are the histograms of the pixelwise forces, fitted to distributions as described in the main text. Red line = Gaussian fit; Blue line = 1^st^ Bimodal fit; Black line = 2^nd^ Bimodal fit. Data points in the graphs represent the average FRET efficiency across each cell measured, and the bar represents the pooled population mean. Comparisons of population means are calculated using one-way ANOVA with Tukey’s correction for multiple comparisons: P-values ≥ 0.123 ns, ≤ 0.0332 (*), ≤ 0.0021 (**), ≤ 0.0002 (***), ≤ 0.0001 (****).

The conformational states of vinculin are proposed to be spatially compartmentalized, with the autoinhibited (closed) state predominantly cytoplasmic and the active (open) state at focal adhesions where it is under tension while bound to talin and actin^49–53^. Talin is a well-established binding partner of vinculin^52–56^, and RIAM and vinculin share a binding site on talin, implying that binding of both proteins is mutually exclusive. However, immunoprecipitated GFP-Vinculin expressed in MEF^Vinc-/-^ formed a complex with both RIAM and talin (Figure 2b). This demonstrates that these proteins are in sufficient proximity to be isolated biochemically from cells, and that vinculin and RIAM may form a complex that is distinct from that with talin. Moreover, levels of RIAM – but not talin - were reduced in this complex following treatment of cells with the microtubule disrupting agent nocodazole (Figure 2b) which also stabilizes focal adhesions^57^. This is consistent with previous reports of lower RIAM levels in mature FA^38^, and suggests reduction of the RIAM-vinculin complex in favor of talin-vinculin binding.

To further investigate the RIAM-vinculin interaction, FRET efficiency distributions were obtained from histograms of the intensity-weighted lifetimes of cells expressing EGFP-vinculin with or without the co-expression of RIAM N- or C- terminally tagged with mScarlet-I (Figure 2c). As expected, co-expression of mScarlet-tagged RIAM led to increased FRET efficiency; equivalently a reduced EGFP fluorescence lifetime (Supplementary Figure S12). Notably, a higher FRET efficiency (16.4 ± 0.4) was observed for N-terminally tagged RIAM compared to C-terminal (13.5 ± 0.7) consistent with previous *in vitro* evidence^38^ and supporting the AlphaFold3 predictions (Figure 2a) that the interaction occurs between the N-termini of both proteins.

The dependence of the RIAM-vinculin complex on FA maturation state indicated a potential role for tension-dependent vinculin conformation in the control of RIAM binding. To address this, we leveraged a previously developed vinculin intramolecular tension sensor (vincTS) comprising of a mTFP1 donor fluorophore and mVenus acceptor fluorophore linked by a 40-amino acid flagelliform linker (TSMod) inserted into full-length vinculin ^36,37,58^. Vinculin adopts an open conformation once bound to both talin and F-actin and under tension^26,36,49,50,55,59^. The same tension sensing module lacking the C-terminal actin-binding domain (vincTL) was used as a high FRET, no tension, control. High variation in mean FRET efficiencies was seen in both FA containing vincTS and those containing vincTL between different cells (Figure 2d and supplementary Figure S13). Previous experiments using the same tension sensor demonstrated static FA with an average force of ≈ 2.5 pN^36^. As individual vinculin proteins bind within adhesions and diffuse in and out of the FA complex, we expect a full range of possible FRET values. The average force applied per pixel can be calculated from^36,37^:

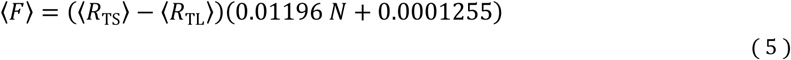

Where *R* _TS_ and *R* _TL_ are the calculated separation distances for the vincTS and vinculin-TL constructs, respectively, and *N* is the number of amino acids in the linker. Both *in vitro* and *in vivo* studies^36,37,43,58^ utilizing the TSMod-based sensor have shown that the 40 amino acid linker used in the vincTS and vincTL constructs is elastic and has an intracellular compliance of approximately 0.478 nm·pN^-1^. From these FRET data, it is possible to calculate from Eq. (5) an average intracellular tensile force ⟨F⟩ = 3.0 ± 0.3 pN. A representative tension map for a cell expressing the VincTS construct and fitted with a bimodal Gaussian distribution illustrates two populations with different levels of tension across vinculin in FAs. Low and high intramolecular tension across FL vinculin was 𝑥_1_ = 2.7 ± 0.4 and 𝑥_2_ = 3.2 ± 0.2 pN respectively (Figure 2d and Table 3).

**Table 3.** Comparison of Bimodal fitting parameters for VincTS and VincTS + mScarlet-RIAM using the Cascade FRET energy transfer model. A summary table detailing the lifetime. Standard deviation and fractional intensity for each peak in the bimodal modal for lifetime and force for VincTS and for VincTS + mScarlet-RIAM. Where 𝑥 = the mean; 𝜎 = standard deviation, and A = amplitude of the bimodal Gaussians.

| Lifetime and force bimodal fit parameters for VincTS only |  |  | Lifetime and force bimodal fit parameters for VincTS + mScarlet-RIAM |  |  |
| --- | --- | --- | --- | --- | --- |
| Lifetime Bimodal Fit Parameters: |  |  | Lifetime Bimodal Fit Parameters: |  |  |
|  | Lifetime (ns) | Std. dev. |  | Lifetime (ns) | Std. dev. |
| $x_1$ | 1.893 | 0.039 | $x_1$ | 2.154 | 0.182 |
| $\sigma_1$ | 0.249 | 0.026 | $\sigma_1$ | 0.437 | 0.121 |
| $A_1$ | 0.838 | 0.080 | $A_1$ | 0.584 | 0.993 |
| $x_2$ | 2.547 | 0.072 | $x_2$ | 3.146 | 0.270 |
| $\sigma_2$ | 0.305 | 0.051 | $\sigma_2$ | 0.406 | 0.181 |
| $A_2$ | 0.571 | 0.048 | $A_2$ | 0.362 | 0.118 |
| Forces Bimodal Fit Parameters: |  |  | Forces Bimodal Fit Parameters: |  |  |
|  | Force (pN) | Std. dev. |  | Force (pN) | Std. dev. |
| $x_1$ | 2.737 | 0.080 | $x_1$ | 3.067 | 0.0583 |
| $\sigma_1$ | 0.413 | 0.046 | $\sigma_1$ | 0.409 | 0.0394 |
| $A_1$ | 0.678 | 0.065 | $A_1$ | 0.784 | 0.0666 |
| $x_2$ | 3.172 | 0.019 | $x_2$ | 3.454 | 0.0219 |
| $\sigma_2$ | 0.161 | 0.030 | $\sigma_2$ | 0.114 | 0.0292 |
| $A_2$ | 0.686 | 0.140 | $A_2$ | 0.633 | 0.1345 |

To investigate RIAM-vinculin interactions as a function of force across vinculin, vincTS was expressed ± mScarlet-RIAM in MEF^Vinc-/-^ cells for analysis using the developed FRET cascade (Figure 3a). mTFP1- vinculin + mScarlet-RIAM exhibited an average FRET efficiency of 20.0 ± 3.0% (Figure 3b), similar to that seen with GFP-vinculin (Figure 2C). Co-expressing mScarlet-RIAM and vincTS significantly increased the average FRET efficiency from 12.3 ± 6.0 % to 29.1 ± 8.8% (Figure 3b). However, FRET efficiency of vincTL also increased from 17.0 ± 5.7% to 28.9 ± 3.3% with the addition of mScarlet-RIAM (supplementary Figure S14), indicating FRET between mTFP1 and mScarlet can occur in the presence and absence of vinculin-F-actin binding. Further analysis in cells treated with the ROCK inhibitor (H1152) to prevent actomyosin contractility showed no change in FRET efficiency of the RIAM- vinculin interaction (supplementary figure S15). This would imply that RIAM binds to vinculin in a force-independent manner.

**Figure 3.**
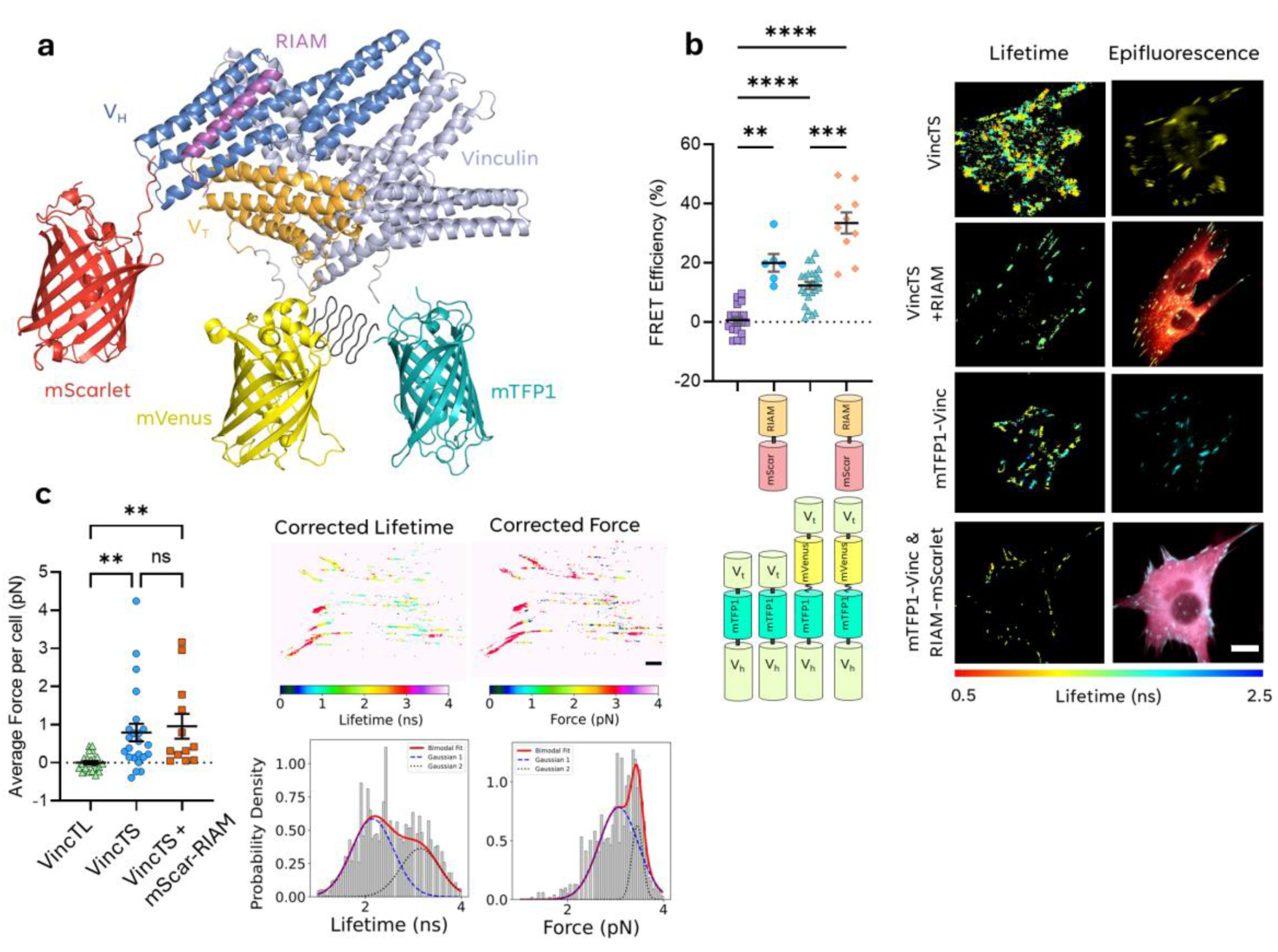
Three-color FRET model applied to the vinculin tension sensing biosensor with mScarlet-RIAM in fixed MEF^Vinc-/-^ cells. Shows a model of the VincTS tension sensor in the presence of mScarlet-RIAM (1-27 aa). This model was created in Pymol using a blend of PDBs (7PNN, mVenus; 4Q9W, mTurquoise2; 5LK4, mScarlet-I) and AlphaFold3 models, such as the previously described Vinculin-RIAM model and the spider-silk linker (GPGGA)_8_. b) Fluorescence lifetime data for the vincTS with mScarlet-RIAM c) The corrected force per cell for vincTS ± mScarlet-RIAM transfected MEF^Vinc-/-^ cells. Data points are the average force across the cell, with the bars representing the pooled population mean. To the right is a corrected lifetime and corrected force map of a typical cell expressing vincTS + mScarlet-RIAM with the corresponding histograms of lifetime and force shown below with fitted curves (as described in the main text). Red line = Gaussian fit; Blue line = 1^st^ Bimodal fit; Black line = 2^nd^ Bimodal fit. c), d), Population means are compared by one-way ANOVA with Tukey’s correction for multiple comparisons: P-values ≥ 0.123 ns, ≤ 0.0332 (*), ≤ 0.0021 (**), ≤ 0.0002 (***), ≤ 0.0001 (****).

To determine if VincTS was under tension when mScarlet-RIAM was present, the energy transfer rate contribution between mTFP1 and mScarlet was calculated and substituted into Eqn 3 to calculate FRET rate independent of mScarlet-RIAM. This allowed us to determine the tension levels across VincTS in the presence of mScarlet-RIAM and compare it to the levels of tension when mScarlet-RIAM is absent. The tensile force for a typical single cell expressing vincTS + mScarlet-RIAM was found to be ⟨F⟩ = 3.2 ± 0.3 pN (standard deviation). This value was determined from a histogram of all the pixelwise values as shown in the tension map in Figure 3C. Moreover, when a bimodal fitting model is applied to the force histogram, two distinct peak forces are identified, suggesting the presence of two sub- populations: x_1_ = 3.1 ± 0.4 pN and x_2_ = 3.5 ± 0.1 pN (Figure 3C and Table 3).

## Discussion

In this study, we developed and validated a three-color Cascade-FRET system capable of quantifying complex molecular interactions and dynamics *in vitro* and in cells. Our methodology expands pair-wise FRET by integrating an additional energy transfer step, providing a more nuanced approach for elucidating the spatial organization of biomolecular complexes. We applied this system to study the interaction of vinculin, a mechanosensitive focal adhesion protein, with its putative binding partner, RIAM, under tension.

The results obtained from purified protein constructs confirm the predictive capabilities of the Cascade-FRET system as these results recapitulate an Alphafold3-predicted structure. Discrepancies in the predicted separation distances are accounted for in the experimental error and likely structure relaxation due to solvation^46^. The results demonstrate that measurements of FRET transition rates can be used to verify structures and distances using a robust, straightforward, theoretical framework. This represents a novel approach to the analysis of structure-function relationships.

We suggest that FRET measurements can be used to determine unknown structures which correlate with a predictive model (Alphafold3). Distances between FRET pairs were determined using FRET efficiencies and Förster radii (Supplementary Tables 1 & 2). A model of the three-color mTurq2- mVenus-mScarlet-I protein predicted shorter distances between fluorophores compared to those measured from the energy transfer rates and determined by negative stain TEM (Supplementary Table S3). The separation distances are all in general agreement, describing a trigonal planar arrangement for the three-color FRET cascade model protein. To fully determine FRET transfer rates within the structure, we introduced a novel G68A mutation in mVenus to produce a non-fluorescent and non-absorbing variant as a FRET control without significantly altering the structure of mVenus (Supplementary Figure S4). This is essential for the characterization of the model system. While other mutations could have been used to create a dark version of mVenus, such as the Q69M and F46L^60,61^, they do not prevent the formation of a functional fluorochrome in the same manner as the G68A mutation.

We successfully demonstrated that vinculin interacts with RIAM in cells, a finding that corroborates previous *in vitro* studies^38^ . Our results provide evidence for the direct binding of RIAM to the N-terminus of vinculin, with higher FRET efficiencies observed for N-terminally tagged RIAM compared to its C-terminally tagged counterpart. This agrees with AlphaFold3 predictions, underscoring the importance of the N-terminus in the interaction.

Using a vinculin tension sensor, we quantified the forces exerted on vinculin within focal adhesions. The average force, 〈F〉, determined by the average FRET efficiency and equation 5, was 3.0 ± 1.0 pN, which is consistent with previous reports^36^ and underscores the heterogeneity of tension in individual adhesions. This heterogeneity likely reflects the dynamic nature of focal adhesions, where vinculin undergoes rapid changes in positioning, F-actin association and binding partners in response to mechanical cues^62–65^. Cascade-FRET allowed us to spatially resolve these forces, revealing how the interaction of vinculin with RIAM correlates with changes in mechanical tension. The average force across vincTS in the presence of mScarlet-RIAM when corrected for the energy transfer between mTFP1 and mScarlet was found to be 3.2 ± 0.3 pN. This value is within the standard error of the vincTS alone where <F< = 2.95 ± 0.97 pN (Figure 3C). Furthermore, when comparing the average tensile forces across each cell for the vincTL, vincTS and vincTS + mScar-RIAM we report that there is no significant difference in average cellular force between the two vincTS conditions (Figure 3C). This indicates that the vinculin-RIAM complex associates when there is moderate force acting on vinculin. Such insights are crucial for understanding how focal adhesions adapt to mechanical stress and how vinculin acts as a molecular clutch to transduce forces between the ECM and the cytoskeleton^49,65–67^.

The data presented in this manuscript has culminated in an updated model of vinculin recruitment through its interaction with RIAM (Figure 4). We have shown vinculin and RIAM interact in cells, supporting the *in vitro* interaction demonstrated previously^38^. Once RIAM associates with the auto-inhibited conformation of vinculin, much like RIAM associates with talin, the complex relocates to the plasma membrane (Figure 4). Vinculin may associate with the R8 domain of talin in the absence of mechanical force^49,59^ and force-induced re-modeling of talin is required for vinculin binding to the R2/3 domains of talin^49^.

**Figure 4.**
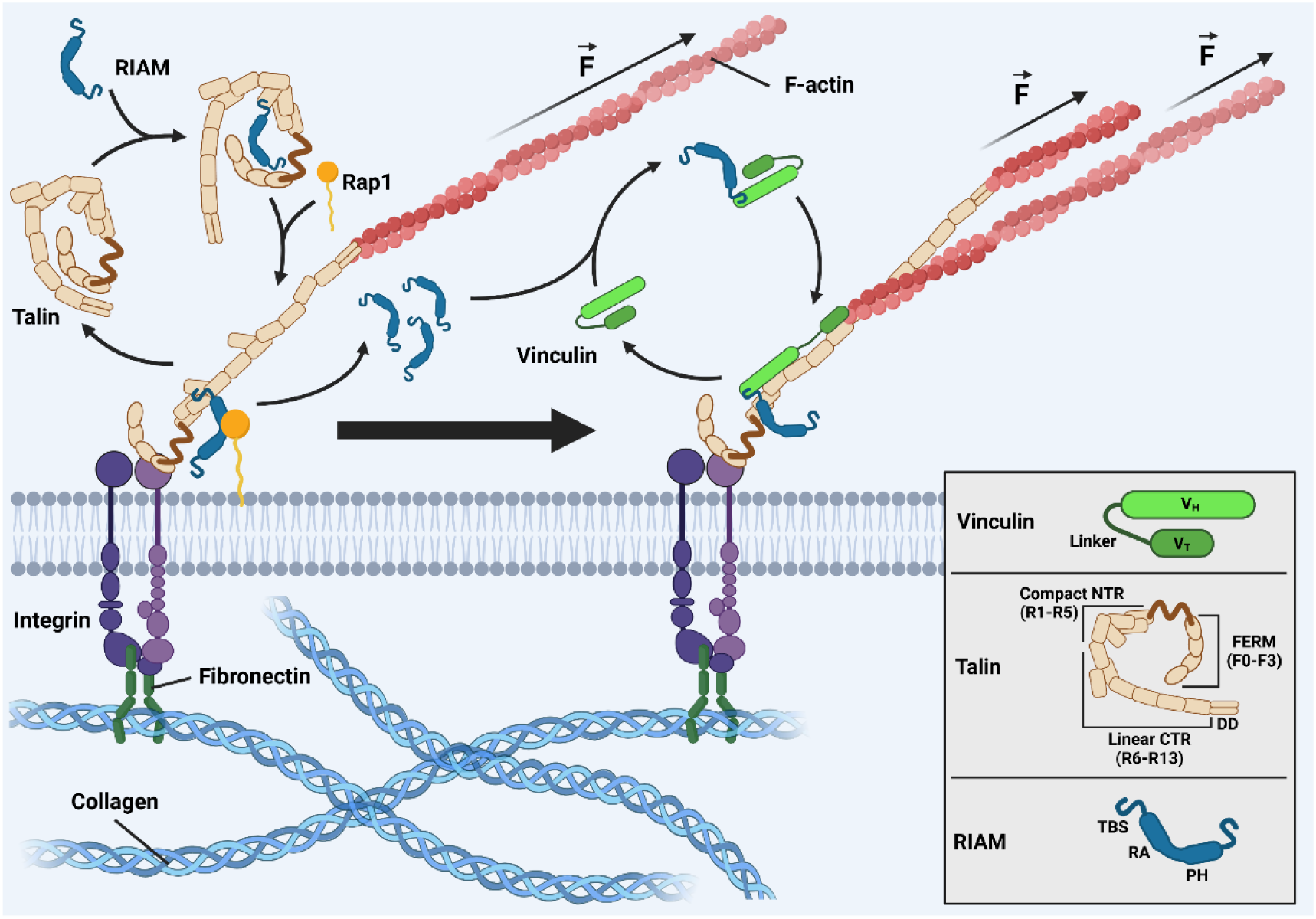
A schematic detailing the key events in our proposed model of vinculin recruitment by RIAM. This scheme illustrates the interaction between the vinculin and RIAM and the activation of vinculin through the binding of talin and f-actin. It also demonstrates how the proposed recruitment of vinculin and talin by RIAM could work. 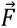 = Tensile force; NTR = N-terminal region; CTR = C-terminal region; DD = dimerization domain. Thin arrows represent the movement of proteins; the thick block arrow represents the evolution of the FA maturation process. The figure was made using BioRender®.

Our data suggests that RIAM mediates vinculin recruitment, as mScarlet-RIAM binds to the vincTS when that construct reports a high average FRET (low tension) (Figure 3b). The lack of change in FRET between GFP-vinculin and mScarlet-RIAM when adding ROCK inhibitor (H-1152) also suggests that RIAM-vinculin binding may not require forces generated from actomyosin (Supplementary Figure S15). We propose that RIAM may dissociate from vinculin, then bind to talin and be further activated through a force-dependent mechanism^38,49^ (Figure 4). We have shown evidence of a RIAM-vinculin interaction in FAs. This could suggest that RIAM plays a role in vinculin recruitment and translocation.

The Cascade-FRET approach offers several advantages over pair-wise FRET methods, including the ability to dissect complex, multi-component interactions with high precision. This methodology can be readily adapted to study other dynamic protein complexes, such as those involved in signal transduction, cytoskeletal organization, or transcriptional regulation.

Future studies could expand the system to include additional fluorophores, enabling the exploration of even more complex biological networks. Moreover, integrating Cascade-FRET with advanced imaging techniques, such as super-resolution microscopy, could provide unprecedented insights into molecular interactions at the nanoscale. Improvements in computational modeling, such as incorporating Amber relaxation into structural predictions, could further improve the agreement of the AlphaFold model with FRET and TEM-based distance measurements.

In summary, our three-color Cascade-FRET system provides a powerful tool for investigating dynamic molecular interactions in vitro and in cells. By applying this methodology to vinculin and RIAM, we have uncovered new insights into the molecular mechanisms of focal adhesion assembly and mechanotransduction. These findings validate the Cascade-FRET system and highlight its potential for broad applications in cell biology and beyond.

## Materials and Methods

### Cell culture & Lipofectamine 3000 Transfection

Vinculin null, Mouse Embryonic Fibroblasts (MEFs) were cultured in high glucose DMEM (Merck, D6429) supplemented with 10% Fetal Bovine Serum (FBS) (Gibco, A4766801), 1% Penicillin-Streptomycin (Thermo Fisher, 15140122), 2 mM L-Glutamine (Thermo Fisher, 25030081) and 1x MEM non-essential amino acids (Thermo Fisher, 11140050), referred to hereafter as complete growth culture medium. Cells were incubated at 37°C with 5% CO_2_. MEFs were seeded at 20,000 cells per 35 mm, on fibronectin coated glass-bottomed µDish (Ibidi, 81158). Plasmid DNA was transiently transfected using Lipofectamine 3000 in a 2:1 DNA:Lipofectamine ratio, (Thermo Fisher, L3000001) per dish.

### Cloning & Site-Directed Mutagenesis

The three-color FRET mTurq2-mVenus-mScarlet-I construct was initially designed using SnapGene® and was made by VectorBuilder® on a proprietary mammalian expression vector. The different FP pairs and single FP constructs were constructed through a series of digestions with restriction endonucleases, which were used to excise specific FPs from the parent three-color construct. The resulting linearized DNAs were then ligated together with T4 DNA ligase and transformed into chemically competent *E. coli* DH5-α cells (Thermo Fisher, 18258012) before plasmid purification using a HiSpeed® ® Plasmid Midi purification kit (Qiagen, 12643). A separate set of constructs were required for bacterial expression. This was achieved by PCR amplification of the required FPs, which were first gel purified and then ligated into a pET151 directional TOPO^TM^ Expression system. Site-directed mutagenesis (SDM) introduced a single point mutation into the mVenus fluorescent protein of the three-color construct at amino acid 67, glycine, which was mutated to an alanine. A pair of primers were designed containing a two-base-pair mismatch in the center of the primer pair. A PCR reaction was then carried out using the Q5® High-Fidelity DNA polymerase (NEB, M0492L), which produced a linearized form of the double-stranded parental template. The linear plasmid was treated with KLD (Kinase, Ligase, and DpnI) mix as part of the SDM kit (NEB, E0554S).

### Protein purification

The fluorescent protein constructs were expressed in the *E. coli* BL21 (DE3) strain, where single colonies were picked and grown overnight at 37°C in 10 mL of LB media before they were used to inoculate 250 mL of auto-induction media^67^ and grown at 18°C for 72-hours. Cells were harvested by centrifugation at 10,000 g for 20 min at 4°C and resuspended in lysis buffer: 50 mM Tris-Cl adjusted to pH 8.0, with 150 mM NaCl, 20 mM Imidazole and protease inhibitor cocktail (Roche) and 5 units/mL of Benzonase endonuclease (Merck). Cell Lysis was achieved by sonication (30% power for a total sonication time of 1 minute and 30 seconds) using a Sonics Vibra-cell VC 750 sonicator. The resulting cell suspension was centrifuged at 18,750 g for 45 minutes at 4°C. The supernatant was removed and micro-filtered through a 0.22 µm pore disc filter (Millipore) before loading on a 1 mL HisTrap column (GE Healthcare) pre-equilibrated with lysis buffer for immobilized metal-ion affinity chromatography (IMAC). Fluorescent proteins were eluted using an imidazole gradient run on an Äkta Pure FPLC chromatography system.

Fractions containing the fluorescent proteins of interest were collected, pooled, and dialyzed overnight at 4°C in an imidazole-free lysis buffer. The Hexa-Histidine purification tag was cleaved by incubating the dialyzed sample for approximately 8 h at room temperature in the presence of the Tobacco Etch Virus (TEV) protease. Once the tag was removed, the fluorescent protein samples were again filtered through a 0.22 µm pore disc filter before re-loading onto the same HisTrap column (GE Healthcare), which was again pre-equilibrated with lysis buffer. The untagged proteins flowed through the column at a flow rate of 5 mL/min and were then collected in the flow-through. The untagged proteins were concentrated using a Pierce™ 10K MWCO Protein Concentrator, before size exclusion chromatography (SEC) on a 16/60 HiLoad Superdex 75 column (GE Healthcare) equilibrated with 50 mM Tris-HCl, pH 8.0, and 150 mM NaCl adjusted to pH 8.0.

### Multiphoton TCSPC FLIM and Analysis

Transfected cells were cultured on borosilicate glass coverslips (VWR, Thickness No. 1.5) coated with fibronectin and fixed in 4% PFA-PHEM solution for 30 minutes at room temperature, 48 hours post-transfection. Subsequently, the cells were permeabilized with 0.2% Triton-X in 50 mM Tris Buffered Saline (TBS) for 20 minutes at room temperature and then quenched with 1 mg/mL NaBH_4_ (also in 50 mM TBS) for a further 20 minutes at room temperature. The Multiphoton-FLIM TCSPC imaging system is a custom system constructed around a Nikon Eclipse Ti-E microscope. This was fitted with a 40x 1.30 NA Nikon Plan-Fluor oil objective and an 80 MHz Ti:Sapphire laser (Chameleon Vision II, Coherent) tuned to 875 or 950 nm for two-photon excitation of mTurquoise2 or mVenus respectively. Photons were collected using a 480/30 nm emission filter for mTurquoise2 or a 525/25 nm emission filter for mVenus (all from SemrockTM) and an HPM 100-40 hybrid detector (Becker & Hickl GmbH). Laser power was adjusted to give average photon counting rates of 10^4^ to 10^5^ photons s^-1^, with peak rates approaching 10^6^ photons s^-1^. Acquisition times of 300 seconds at low excitation power were used to achieve sufficient photon statistics for fitting while avoiding either pulse pile-up^3,68^ or significant photobleaching. All FLIM data were analyzed using a time-resolved image analysis package, TRI2^6^, and were fitted with either a mono-exponential or biexponential decay curve using the Levenberg-Marquardt algorithm. Lifetime data processed in TRI2 produced histograms of pixel frequencies against photon arrival times for every FP imaged in each cell condition and experiment. A custom Python script was written which imported these histograms for each cell in each experiment into a single data frame, and an intensity-weighted average lifetime was calculated for each cell imaged. An average of 10-15 cells per cell condition were imaged in each experiment, and an unweighted average of these (each cell imaged given an equal weighting) was calculated. Mean lifetimes were used to calculate FRET efficiencies and energy transfer rates within the same Python script. Graphs illustrating the spread of average lifetimes and FRET efficiencies, as well as all corresponding statistical tests, were produced using GraphPad Prism 6.

### Force mapping and visualization

A Python script was written to convert lifetime measurements into force measurements within focal adhesions. The histogram of lifetime data per pixel was loaded into the script and a mathematical transformation applied to the lifetime values, resulting in force values. The processed data is then reshaped into a structured format and converted into images—one representing the distribution of forces and another showing molecular lifetimes.

## Acknowledgments

The authors thank Ambrish Kumar for their assistance with protein purification, Mark Pfuhl for providing TEV protease, purification and the Nikon Imaging Centre@ King’s staff for assistance with spinning disc confocal imaging. We acknowledge funding from the UK MRC MR/X012794/1 and Cancer Research UK for Simon M. Ameer-Beg and MRC/EPSRC for the doctoral scholarship of Conor A. Treacy.

## Author Contributions

C.T., M.P. M.A.P and S.A.B. designed the research. C.T., S.P. and T.B. performed research. C.T., T.L.P, M.P, T. B. and S.A.B. wrote the paper.

