## Supplemental Methods and Figures for "Development of three-color FRET measurement of force-dependent sensing of RIAM-Vinculin interactions in focal adhesions"

### Supplementary Methods

**Data analysis.** The energy transfer rate $\Gamma_{DA}$, describes the rate of energy transfer (FRET) from the donor to the acceptor. It can also describe the probability of the donor de-excitation or 'decay' occurring through FRET.[5] Two other decay paths occur on all fluorophores; these are the radiative decay path $\kappa_r$, where the fluorophore emits a photon and the non-radiative decay path, where energy is lost by interactions with the environment, as given by the non-radiative rate $\kappa_{nr}$. There is a reciprocal relationship between fluorescence lifetime and energy transfer rates such that:

$$\frac{1}{\tau_D} = k_r + k_{nr} \quad and \quad \frac{1}{\tau_{DA}} = \Gamma_{DA} + k_r + k_{nr} \quad (1)$$

The two equations in (1) above can be substituted into the classic FRET efficiency equation to give equation 2. From henceforth, the first donor will be known as $D_1$, the first acceptor/second donor as $A_1$and the final acceptor as $A_2$.

$$\langle E \rangle = \frac{\Gamma_{DA}}{\Gamma_{DA} + k_r + k_{nr}} \quad (2)$$

Equation 2 below describes the energy transfer between any given donor-acceptor pair. This can be expanded to provide the equations described in equation 3, which detail the energy transfer rates for all three possible combinations between fluorophores in a three-color system.

$$\langle E_{D_1A_1} \rangle = \frac{\Gamma_{D_1A_1}}{\Gamma_{D_1A_1} + k_r + k_{nr}}$$

$$\langle E_{A_1A_2}\rangle = \frac{\Gamma_{A_1A_2}}{\Gamma_{A_1A_2} + k_r + k_{nr}} \quad (3)$$

$$\langle E_{D_1A_2}\rangle = \frac{\Gamma_{D_1A_2}}{\Gamma_{D_1A_2} + k_r + k_{nr}}$$

$$\langle E_{D_1A_1+D_1A_2}\rangle = \frac{\Gamma_{D_1A_1}}{\Gamma_{D_1A_1} + k_r + k_{nr}} + \frac{\Gamma_{D_1A_2}}{\Gamma_{D_1A_2} + k_r + k_{nr}} \quad (4)$$

As the donor fluorescence lifetimes scale with the FRET efficiencies, the total transfer rate increases as the FRET transfer rate increases proportionally. The time the donor electron spends in the excited state before being emitted decreases[69]. Thus, lifetime measurements provide a valuable tool for elucidating the de-excitation rates acting on the donor. Measured FRET rates are linear and cannot be added to lifetimes as the radiative and nonradiative decay pathways do not scale with increased FRET[69]However, in isolation, we can construct a theoretical average transfer rate and, by extension, FRET efficiency from the measured lifetimes of the donor alone and donor + acceptors 1 and 2. This lifetime should be the reciprocal of the lifetime measured for the construct with one donor and two acceptors (see equation 4).

**Derivation of the force equation.** The relationship between FRET efficiency and distance is shown in Equation 6 below. Where $R_0$ is the Förster radius, the distance between fluorochrome centers where the FERT efficiency is 50%, and $r$ is the separation distance between the fluorochrome centers.

$$\langle E\rangle = 1 - \frac{\tau_{DA}}{\tau_D} = \frac{{R_0}^6}{{R_0}^6 + r^6} \quad (6)$$

Using equation 6, where the $\tau_{DA}$ is the lifetime of mTFP1 in the vincTS construct, and $\tau_D$ is the lifetime of mTFP1 in the mTFP1-vinculin construct. We can rearrange equation 6 to make the separation distance, r, the subject; see equation 7.

$$r = \sqrt[6]{\frac{\tau_D \cdot \tau_{TS}}{1-\tau_{TS}} {R_0}^6} \quad (7)$$

The applied force on any spring can be described by Hook's law (equation 8), Where $k = (0.01196 \cdot N + 0.0001255)36{,}43$, for this instance, N= 40 is the number of amino acids in the nanospring. Resulting in k = 0.478 pN/m. The change in spring distances is given by equation 9. This represents the maximal change in spring length when tensile forces are applied.

$$\Delta x = r_{TS} - r_{TL} \quad (8)$$

$$F = k \cdot \Delta x \quad (9)$$

Combining equations (7), (8) and (9) gives equation (10) below. This fully describes how the change in fluorescence lifetime of the vincTS and vincTL transfected cells results in changes in the average applied tensile force found within focal adhesions.

$$\langle F \rangle = k \cdot R_0 \cdot \sqrt[6]{\tau_D} \cdot \left(\frac{\tau_{TS}}{1-\tau_{TS}} - \frac{\tau_{TL}}{1-\tau_{TL}}\right)^{\frac{1}{6}} \quad \text{simplified as} \quad \langle F \rangle = (r_{TS} - r_{TL}) \cdot 0.478 \text{ pN} \quad (10)$$

**Development of a mutant mVenus with an ablated chromophore.** The chromophores found in all fluorescent proteins comprise only three amino acids; in GFP, these are $Ser_{65}$, $Try_{66}$, and $Gly_{67}$. These amino acids undergo a process of cyclization and dehydrogenation before forming a functional chromophore[70,71] (figure 1c). The $Gly_{67}$ in GFP is highly conserved for many species. FPs isolated from distantly related organisms like *Aequorea victoria* and *Entacmaea quadricolor* possess a glycine in the third position of the three amino acids. Early site-directed mutagenesis work[70,71] a $Gly_{67}$ to Alanine mutation was sufficient to prevent the fully fluorescent chromophore forming in GFP. It was speculated that the cyclization step was impossible due to the steric hindrance induced by the methyl group on the alanine.[71,72] The equivalent mutation in mVenus is $Gly_{68}$ to $Ala_{68}$; this was selected as a possible way to produce a "non-fluorescent beta barrel" or spacer, which could occupy the same physical space of the mVenus[73] but would not have any of the photophysical properties (visible absorption or emission) and, most importantly, would not FRET with the mTurquiose2 donor. The absorption and emission spectra for the purified mTurq2-mVenus$^{G68A}$-mScarlet protein illustrates the absence of an emission peak associated with mVenus at 530 nm, which can be seen in both the unmutated three-color protein and the mVenus only control (figure 1b). This indicates that there is no longer a functional mVenus when the G68A mutation is introduced. Furthermore, we wanted to know if the integrity of the mVenus beta-barrel remained after the introduction of the G68A mutation or if the mutation had caused the beta-barrel to misfold, as previously suggested by Cubitt et al., 1995[71].

The near-UV spectra for the mutated and unmutated mTurq2-mVenus-mScarlet proteins show that there is a large degree of similarity between the two (Supplementary Figure SF4a), with two notable exceptions being the 250-300 nm and 430-580 nm regions. The difference in the latter in the mutant is due to the absence of an active mVenus chromophore, which no longer has an absorbance peak at 515 nm. This can also be seen in the absorbance spectra of the mutated mTurq2-mVenus-mScarlet protein, wherein the absorbance spectra for the same proteins, a difference can be seen for the same region surrounding the 515 nm absorbance peak associated with mVenus: with this also being absent for the mutated mTurq2-mVenus-mScarlet protein. The far-UV spectra (Supplementary Figure SF4c) can reveal important characteristics of the secondary structure of a protein. Electronic transitions associated with the chiral amide groups can be detected with far-UV CD. This is of particular interest, as this can be used to predict the secondary structure of a peptide[74,75]. The three-color fluorescent proteins share very similar far-UV spectra with minimal differences between the proteins mTurq2-mVenus-mScarlet and mTurq2 mVenus$^{G68A}$-mScarlet. The shape of the far-UV CD spectra (Supplementary Figure SF4c) suggests that the proteins are primarily beta-sheets due to the broad trough around 190-220 nm and the rise towards a peak around 195-200 nm. This result confirms that the proteins are formed primarily from beta sheets, as expected, for a pair of proteins comprised of three beta-barrels, each consisting of 11 beta sheets and one coaxial helix per barrel[72,76]. The far-UV Spectra were also recorded over a range of temperatures (Supplementary Figure SF4d (cut-out)); the change in Mean Residue Ellipticity (deg $cm^2$ $dmol^{-1}$) against temperature for a single wavelength at 205 nm was recorded for three-color proteins. A single wavelength was chosen between the two transitional regions to best demonstrate changes in the dihedral angles and, by extension, changes in secondary and tertiary structure caused by temperature. The mVenus-containing protein (red) remains primarily unchanged until approximately 60 °C, whereas the mVenus$^{G68A}$-containing mutant protein (blue) has some significant changes by 50 °C. This suggests that there is a difference in thermal stability between the proteins and that there may still be the possibility that the mutant is not as tightly folded in the absence of a mature chromophore.

The results of secondary structure analysis using[77,78] (Supplementary Figure SF4e) predicted the same secondary structures for the two three-color proteins. This is not surprising, as their far-UV spectra have very little difference. The expected data (green bars) shows some moderate difference but nothing significant, apart from an overestimation of the anti-parallel (right-twisted) beta-sheet and a slight underestimation of the unstructured region. This is a good indication overall that there is not a significant difference between the three-color proteins in terms of the secondary and tertiary structures; the mutation may have reduced the overall stability of that protein, especially at higher temperatures, which may be the reason why previous groups did not successfully purify viable proteins with this mutation.

**Spectrofluorometric & Circular Dichroism measurements.** Excitation and emission spectra for the eluted purified proteins were performed in a 5 mL quartz cuvette, and measurements were taken on a Horiba FluoroMax® 4 Spectrometer. Excitation spectra were recorded between 300 and 600 nm in 1 nm increments with a 5 nm spectral bandwidth. The emission spectra were taken between 400 and 650 nm in 1 nm increments with a 5 nm spectral bandwidth. The same set of excitation and emission wavelengths was used for all constructs. The emission wavelengths were used for the excitation spectra: 480, 530, and 600 nm. For the Emission spectra, the following excitation wavelengths were used: 435, 515, and 590 nm. Ultra-violet and Circular dichroism (CD) spectra of the mTurquoise2-mVenus-mScarlet-I and mTurquoise2-mVenusG68A-mScarlet-I samples were acquired on the Chirascan Plus spectrometer (Applied Photophysics) using a Suprasil rectangular cuvette (Hellma UK & Starna Scientific Ltd). The instrument was flushed continuously with pure evaporated nitrogen throughout the experiment. For the UV-visible spectra, a wavelength range of 230-800 nm was used with a Spectral Bandwidth of 1 nm, a time per point of 0.5 seconds, and a path length of 10 mm. The far-UV CD spectra used a wavelength range of 195-260 nm with a Spectral Bandwidth of 2 nm, a time per point of 1.5 seconds, and a path length of 0.5 mm. Where appropriate, the CD spectra were smoothed with a window factor of four using the Savitzky-Golay filter[79-81]. The far-UV CD spectra of the samples were first recorded at 23°C, cooled to 6◦C then heated to high temperature (94◦C) and then cooled again to 23°C. Multi-wavelength melting profiles monitored between 195 and 260 nm were recorded from 6 to 94◦C during heating. The instrument was equipped with a Quantum TC125 Peltier (NorthWest) set to change the temperature from 6 to 94◦C at 1◦C per minute and 1.5 second time-per-point CD measurement time. The total scan time was 2 minutes per spectrum, and a 1 nm step-size was employed in the 195 to 260 range with a 2 nm Spectral Bandwidth. The temperatures were measured directly with a thermocouple probe in the sample solution, and the buffer baseline was auto-subtracted. Melting temperatures were determined from the derivative CD vs Temperature spectra and fitted using a Levenberg–Marquardt algorithm (LMA) on the Van't Hoff isochore. (Global 3, Global Analysis for T-ramp Version 1.2 built 1786, Applied Photophysics Ltd, 2007-2012).

### Supplementary Figures

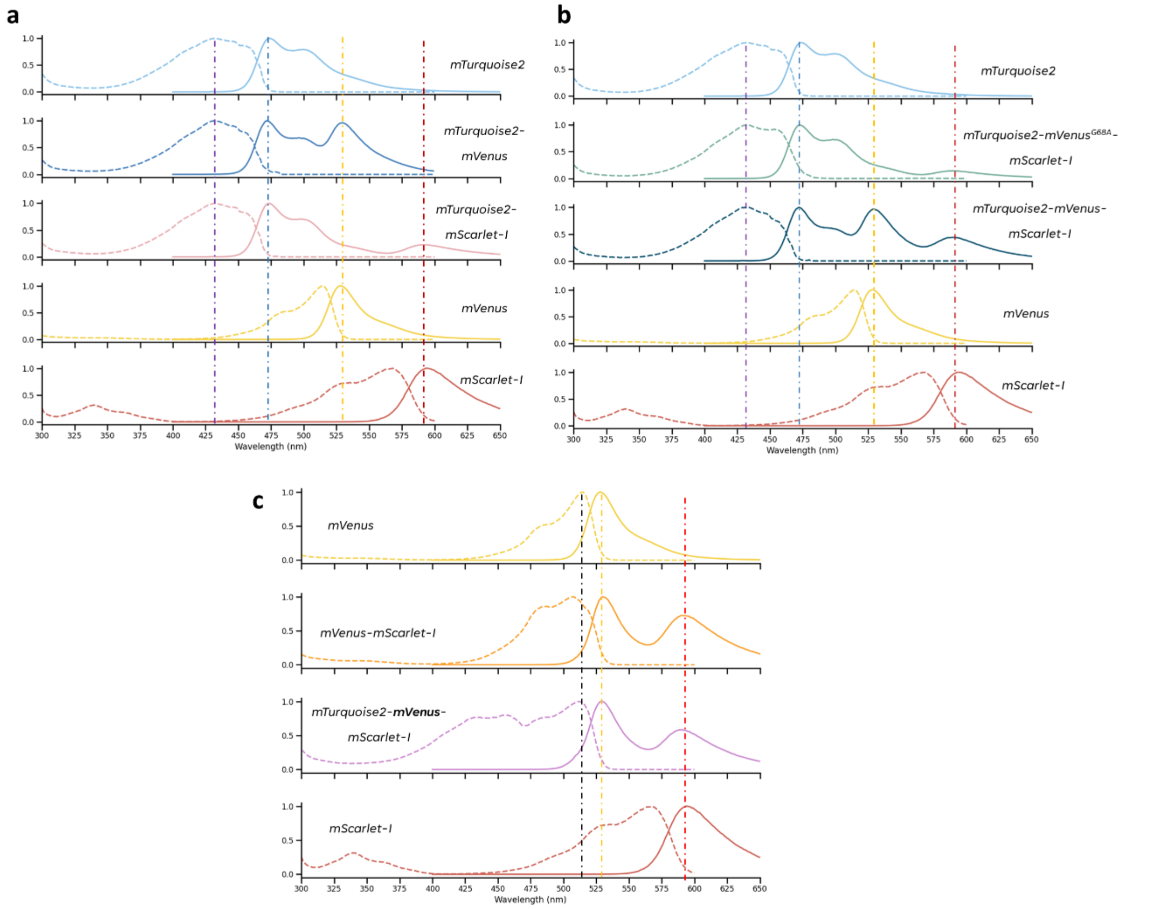


**Fig. S1. Excitation and Emission spectra for fluorescent proteins. Spectroscopic measurements were taken on a FluoroMaxR4 spectrofluorometer (Horiba), and 2 mL of purified fluorescent protein was loaded into the spectrofluorometer in a quartz cuvette before the excitation and emission spectra were measured for each protein. Excitation spectra are in dashed lines, and the emission spectra are in solid lines, and all spectra are normalized to their peak intensities. The emission spectra were collected by exciting the mTurq2 @ 430 nm for Constructs containing mTurq2 and collecting the emission spectra from 450-600 nm. Where mVenus was the donor, excitation @ 510 nm and emission spectra were collected from 530-600 nm.**

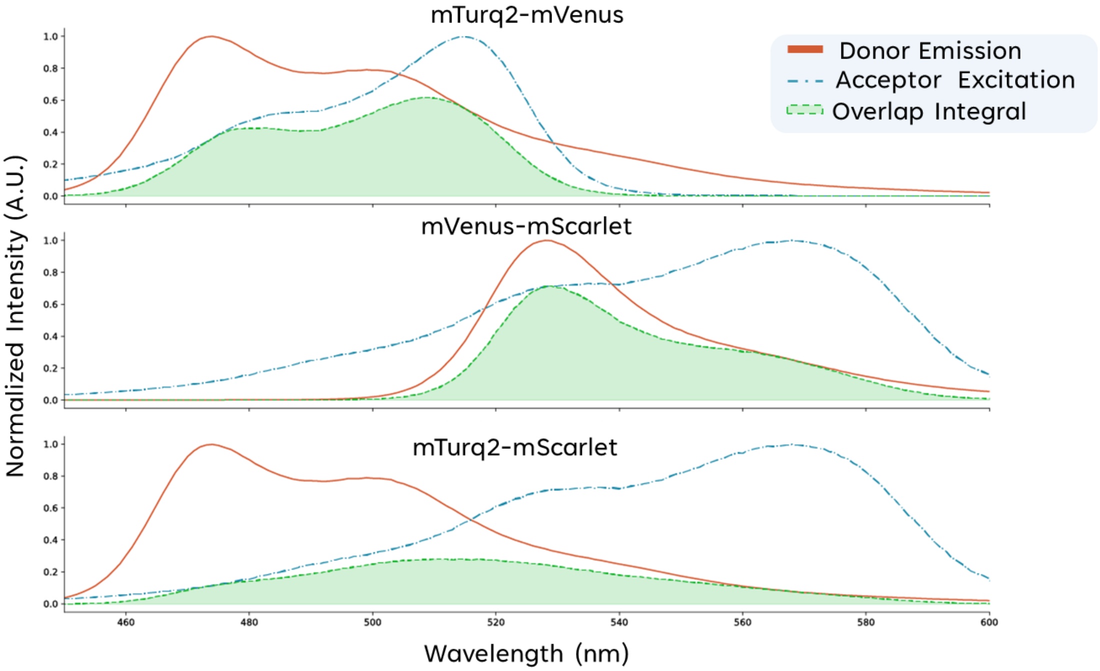


**Fig. S2. Overlap Integrals for the FRET pairs found in the purified FPs**: The donor emission spectra (red solid line), the acceptor excitation spectra (blue dot-dashed line) and the area overlap integral (green shaded area under the green dotted line) that lie between the donor and acceptor. A) the mTurquoise2-mVenus FRET pair, B) the mVenus-mScarlet FRET pair and C) the mTurquoise2-mScarlet FRET pair.

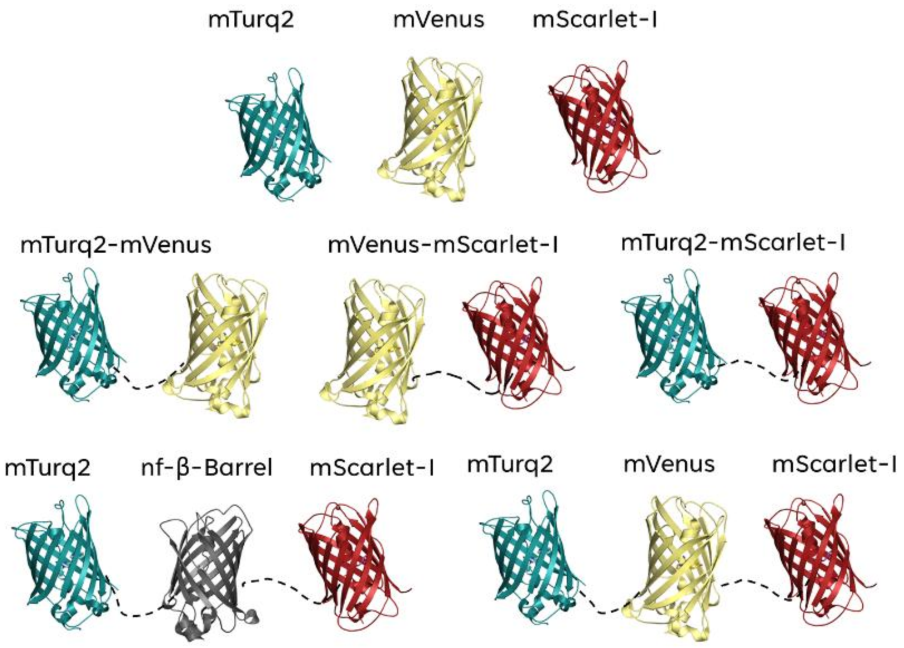


**Fig. S3. Cartoons of the chimeric fluorescent proteins:** Top row: The three individual FPs mTurquoise2, mVenus and mScarlet-I. Middle row: The three twin FPs; mTurquoise2-mVenus, mVenus-mScarlet-I, mTurquoise2-mScarlet I. Bottom row: The two triple FP constructs: The fully functional mTurquoise2-mVenus-mScarlet-I construct and the mutated mTurquoise2-mVenusG68A-mScarlet-I construct. All images of the fluorescent proteins were created from their PDB accession codes and modelled in Pymol. Linkers shown in black are dotted lines for illustrative purposes, and linker sequence GGSGGS is for all twin and triplet constructs.

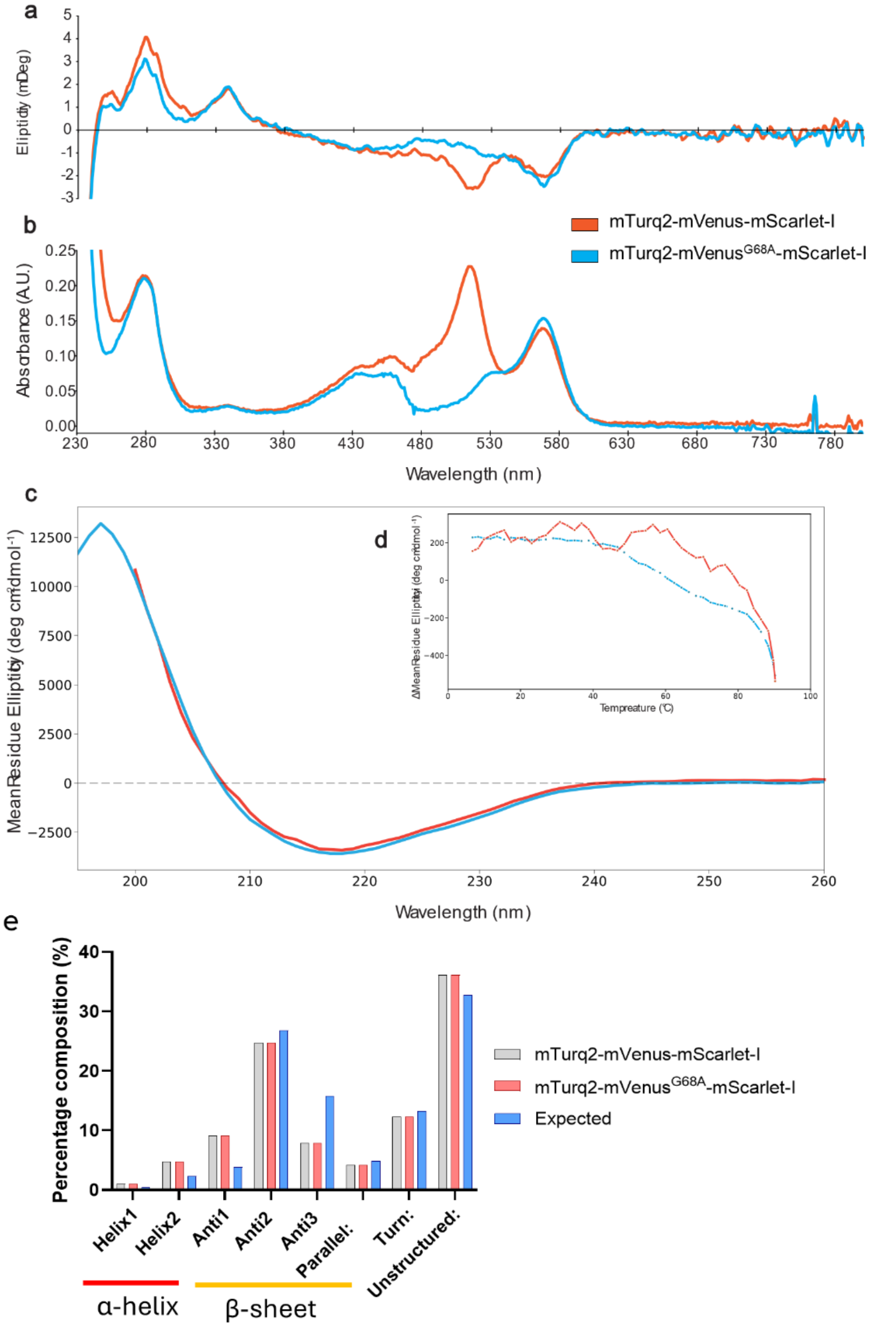


**Fig. S4. Near and Far-UV Circular Dichroism Spectra for the two three-colored FRET Cascade FPs:** a) shows the near-UV CD spectra, b) the near UV-vis absorption spectra for the two three-colored FPs. c) the far-UV CD spectra. d) The change in Mean Residue Ellipticity as a function of heating at a specific wavelength of 205 nm. E) Percentage composition versus secondary structure type for the two three-color FPs and the expected --average secondary structure for the three fluorophores. CD spectra analysis was performed using the BeStSel

(Beta Structure Selection) single spectrum analysis online tool *(https://bestsel.elte.hu/index.php)*.

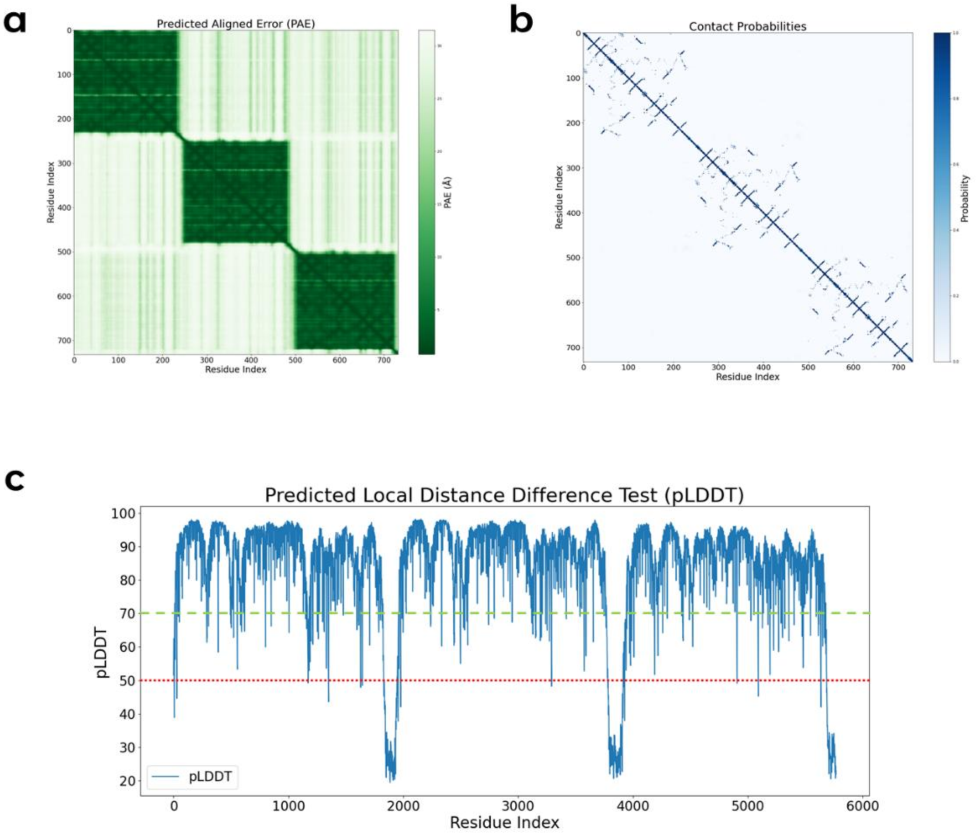


**Fig S5. AlphaFold3 validation on the three-color model**. a) The predicted aligned error (PAE) plot shows dark green areas for the three fluorophores in the three-color cascade model, indicating that each FP is well-defined. Note that the green shading is weaker between FPs, indicating the inter-domain configuration is not as well defined. Indicating more potential flexibility between linked FPs. b) Contact probability analysis suggests that interactions are confined mainly within each fluorescent protein (FP), with weakly conserved contacts reflecting the low-affinity self-association sites. These non-specific interactions likely drive the assembly into a triangular shape. c) The predicted local distance difference test (pLDDT) is a per-residue measure of local confidence and is plotted for each residue of the AlphaFold3 model. There is high confidence (>70) for nearly all residues apart from the two linker regions (amino acids 275-285 and 524-533). Regions with a high pLDDT score (>70) are a green dashed line, and those with a very low pLDDT score (< 50) are indicated with a red dotted line.

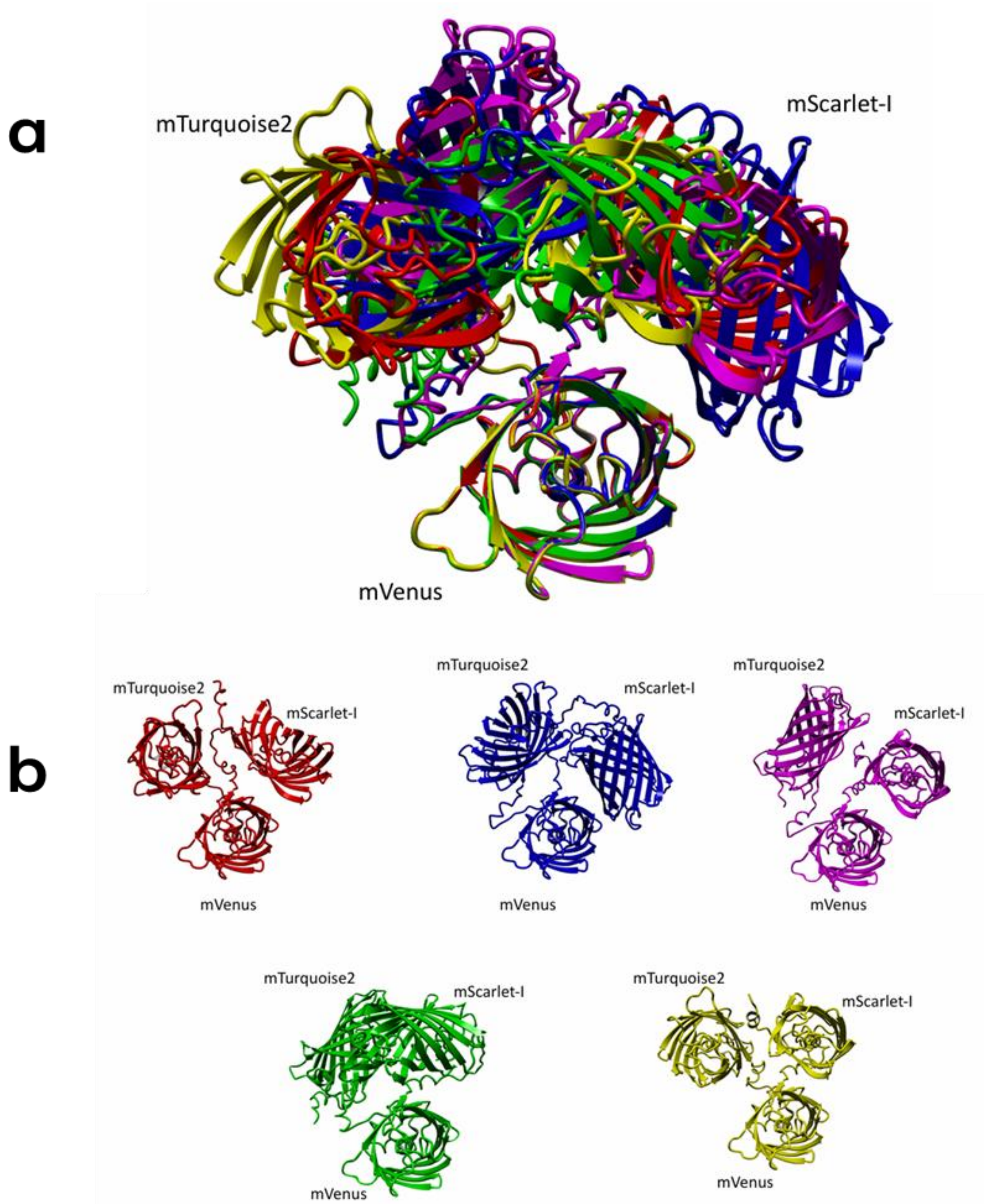


**Fig S6. AlphaFold3 alternate models.** a) Superimposed structures centred about mVenus for the five AF3 models generated. b) individual models generated using AF3.

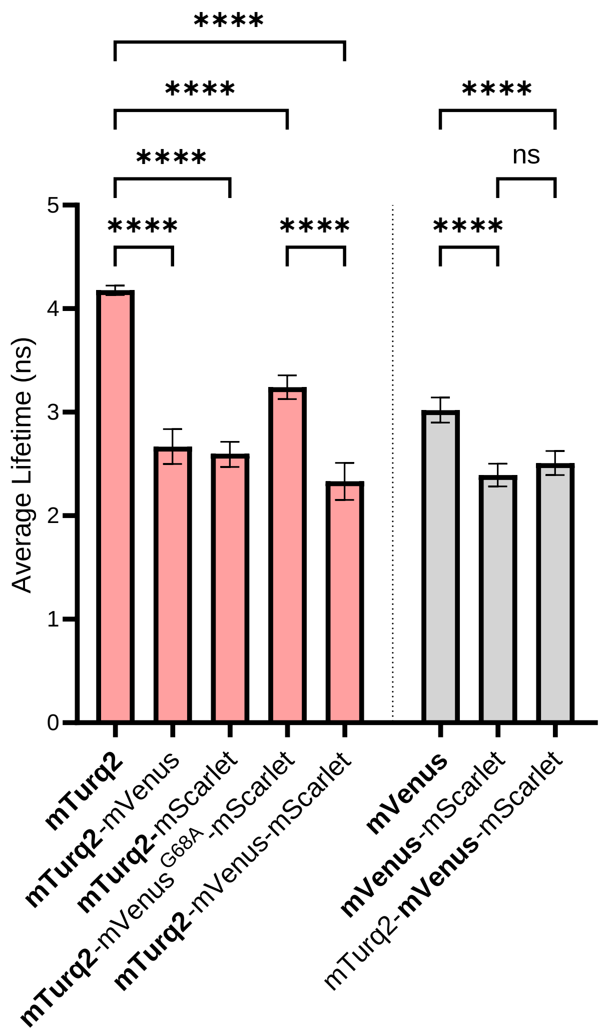


**Fig S7. Average lifetimes for the FRET Cascade:** Fluorescent Proteins Measured in Solution: graph showing the average lifetime recorded for each FP; error bars indicate standard error. N=10 measurements per condition across three separate technical repeats. P-values ≥ 0.123 ns, ≤ 0.0332 (*), ≤ 0.0021 (**), ≤ 0.0002 (***), ≤ 0.0001 (****).

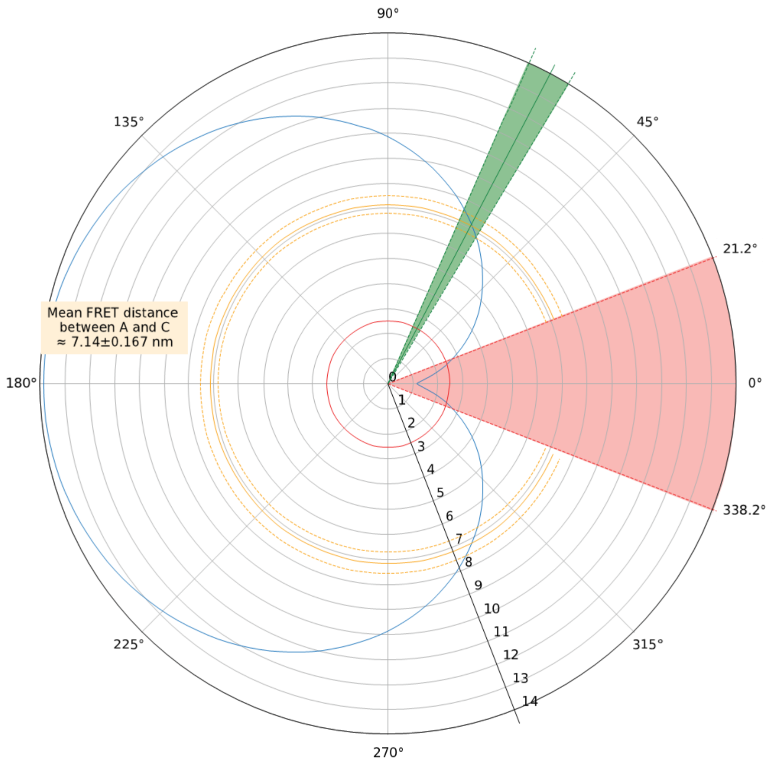


**Fig S8. Polar plot of separation angle versus distance for the mTurq2- mVenus-mScarlet protein:** Shows a polar plot of theta. This separation angle lies between mTurq2 and mScarlet against the distance between mTurq2 and mScarlet.

a

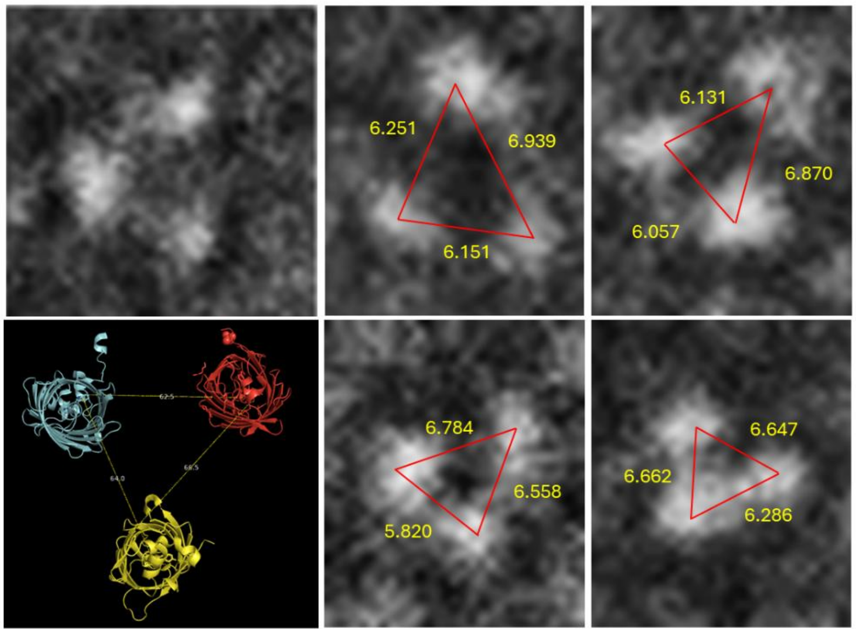


b

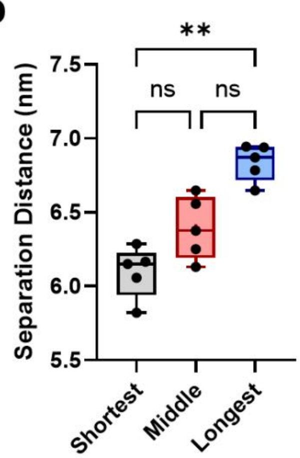


c

| | | Shortest | Middle | Longest |
|---|---|---|---|---|
| TEM Measured | Mean | 6.096 | 6.393 | 6.837 |
| | SEM | 7.80x10-2 | 9.51x10-2 | 5.56x10-2 |
| FRET Calculated | Mean | 6.25 | 6.41 | 6.91 |
| | SEM | 5.49x10-2 | 5.23x10-2 | 1.03x10-1 |
| Difference | | -2.46% | -0.27% | -1.06% |

**Fig. S9. Negative stain TEM data**. Average Separation distance between β-Barrels Determined by Negative Stain TEM: a-e) Negative Stain TEM micrographs acquired at 120kV of the three color mTurq2-mVenus-mScalet-I protein. Distances between beta-barrels measured in Fiji (ImageJ). f) The predicted model is rendered in Pymol, a top-down projection for comparison with TEM data. g) Bar chart of average distances between β-barrels in the TEM micrographs. H) The summary statistics table outlines the mean and standard deviations for the distances measured (TEM micrographs) and calculated (FRET calculations), as well as the percentage difference between the two. N=3 measurements per condition across three separate technical repeats. P-values ≥ 0.123 ns, ≤ 0.0332 (*), ≤ 0.0021 (**), ≤ 0.0002 (***), ≤ 0.0001 (****)

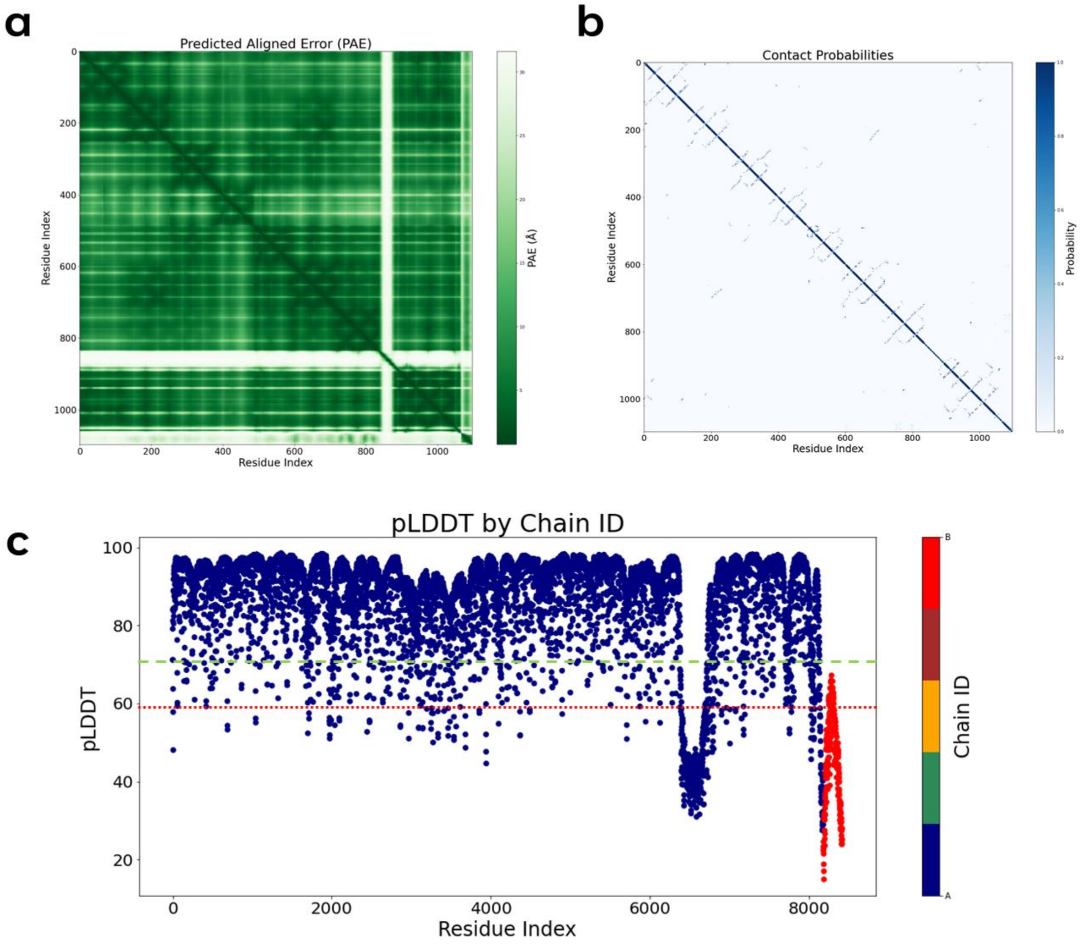


**Fig S10. AlphaFold3 RIAM-vinculin model**. a) The predicted aligned error (PAE) plot shows dark green areas, indicating that the model for the vinculin head domain (residues 1-885) and vinculin tail domains (residues 1401-1586) are highly aligned with low error. There is a notable gap for the vinculin neck region, which is highly dynamic. Significant contact probabilities are shown for RIAM interacting with vinculin b) Contact probabilities show the intradomain interaction within the vinculin molecule. c) The predicted local distance difference test (pLDDT) is a per-residue measure of local confidence and is plotted for each residue of the AlphaFold3 model. There is high confidence (>70) for nearly all residues in the vinculin molecule, apart from the linker neck region. RIAM (red) is also plotted and shows a generally low pLDDT score. Regions with a high pLDDT score (>70) are a green dashed line, and those with a very low pLDDT score (< 50) are indicated with a red dotted line.

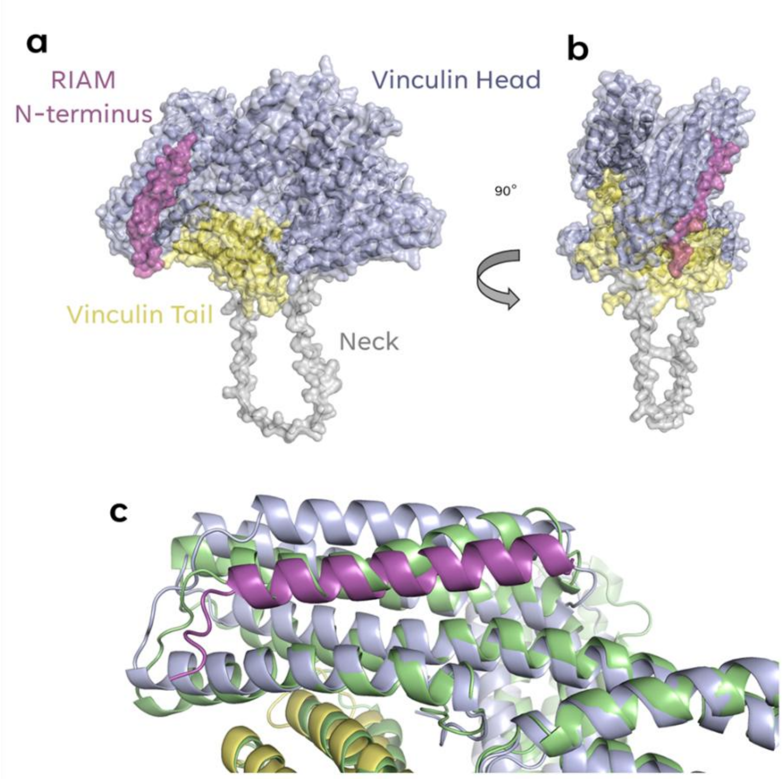


**Fig. S11. Vinculin autoinhibited conformation**. a) Shows auto-inhibited vinculin head domain bound to the vinculin tail domain with RIAM bound to the D1 sub-domain of the vinculin head domain. Image produced in Pymol from AlphaFold3 model. b) a 90-degree rotation of the RIAM-vinculin model. c) A diagram illustrating the re-positioning of vinculin $V_H$ D1 alpha helices in the presence (blue) and absence (green) of RIAM (purple).

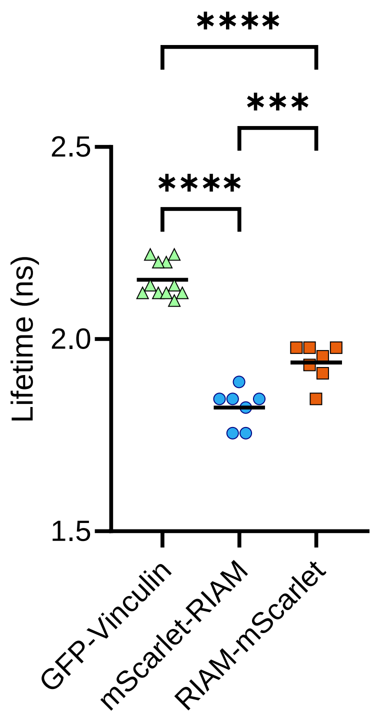


**Fig. S12.Fluorescent lifetime data for the RIAM-vinculin interaction determined by TCSPC-FLIM:** Average lifetime data per construct. Significance was determined through a one-way ANOVA with Tukey corrections for multiple tests. N= 10 cells imaged per condition, with P-values ≥ 0.123 ns, ≤ 0.0332 (*), ≤ 0.0021 (**), ≤ 0.0002 (***), ≤ 0.0001 (****)

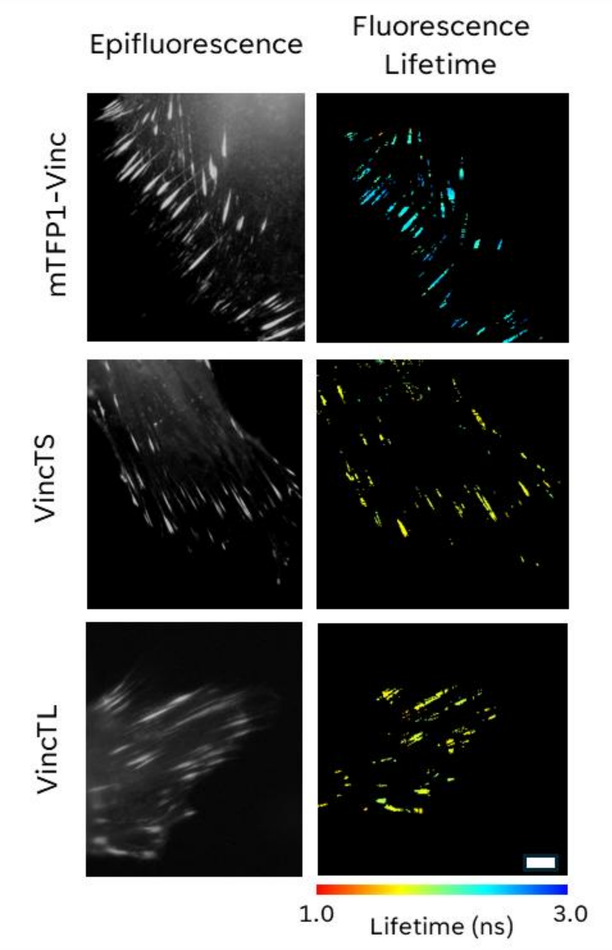


**Fig. S13. The Vinculin Tension Sensor determined by TCSPC-FLIM** shows epifluorescence and lifetime images of MEF$^{vinc-/-}$ transfected with mTFP1-vinculin (donor only), vincTL or vincTS. Scale bar = 5 µm.

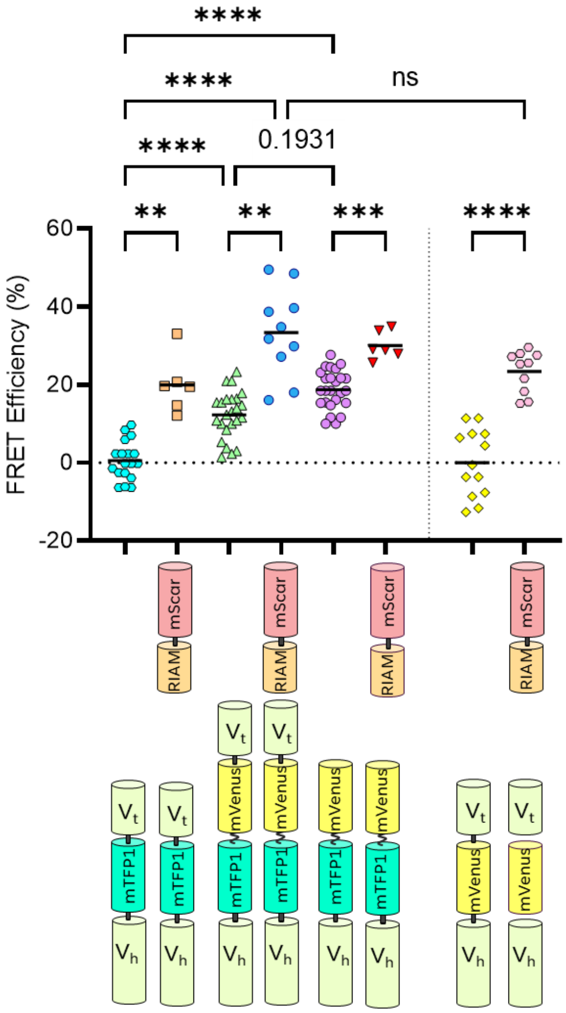


**Fig.S14: Fluorescence Lifetime Distributions for vinculin-TS with mScarlet-RIAM in Fixed cells:** a) Summed average lifetime plots for the various Vinc-TS & mScarlet-RIAM conditions along with Vinc-TS alone and Teal-vinculin alone lifetime plots. b) Plots of normalized pixel frequency against lifetime for the three masked Vinc-TS & mScarlet-RIAM conditions. P-values ≥ 0.123 ns, ≤ 0.0332 (*), ≤ 0.0021 (**), ≤ 0.0002 (***), ≤ 0.0001 (****)

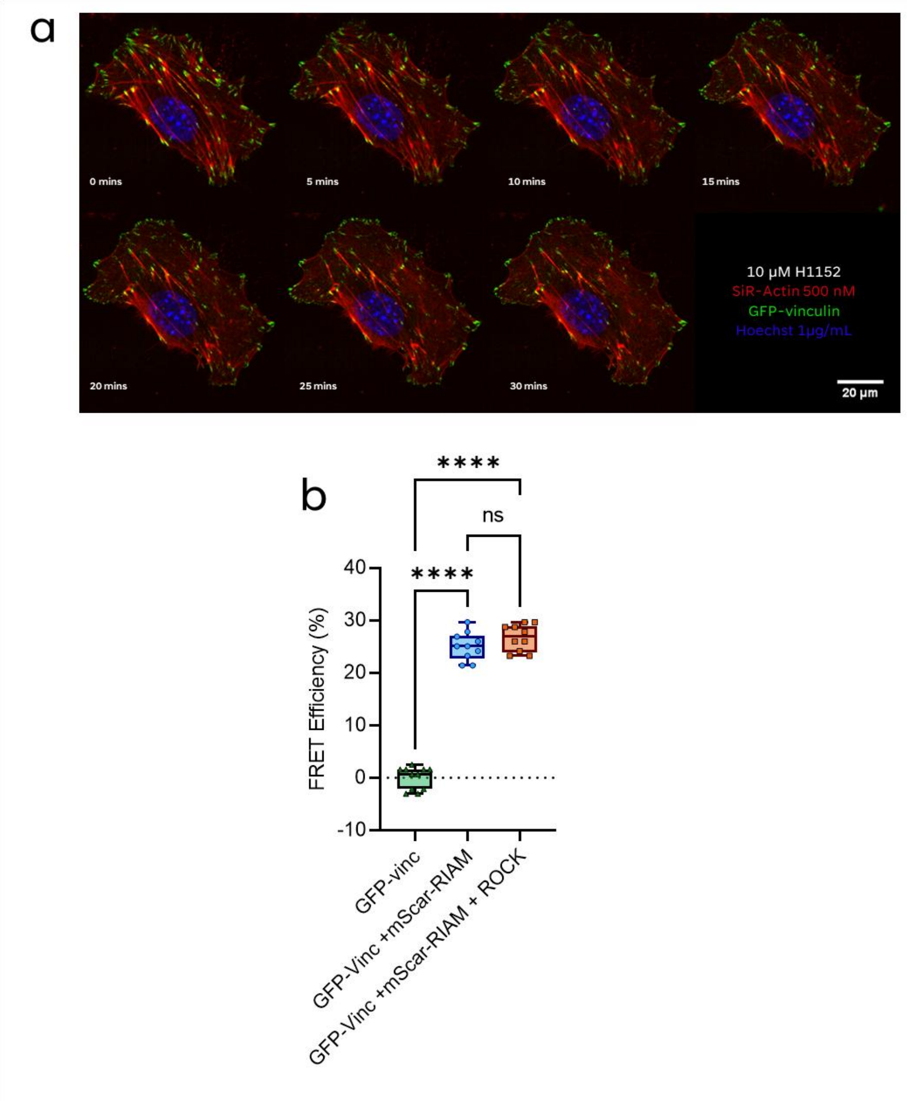


**Fig. S15: Live cell staining of cytoskeletal and focal adhesion proteins in the presence of ROCK Inhibitor H-1152:** a) MEFs transfected with GFP-vinculin and stained either with SiR-Actin at 500 nM, and Hoechst nuclear stain at 1 µg/mL. Cells were treated with 10 µM ROCK inhibitor (H-1152) for 30 minutes during live-cell imaging. Imaging was conducted on a Spinning Disc Confocal imaging system (NIC@King's). b) Box and whisker plot of the average FRET efficiencies per construct. Significance was determined through a one-way ANOVA with Tukey corrections for multiple tests. N= 10 cells imaged per condition, with P-values ≥ 0.123 ns, ≤ 0.0332 (*), ≤ 0.0021 (**), ≤ 0.0002 (***), ≤ 0.0001 (****).

### *Supplementary Tables*

**Table S1. FP summary table:** A table detailing the Quantum yield of the donor (QYD), the Förster radius of the FRET pair in nm, and the overlap integral (Jλ) in $M^{-1}$ $cm^{-1}$ $nm^4$

| | mTurq2 → mVenus | mTurq2 → mScarlet-I | mVenus → mScarlet-I |
|---|---|---|---|
| Energy Transfer Rates, r (nm) | 6.41 | 6.25 | 6.91 |
| AlphaFold3 model, r (nm) | 4.04 | 4.32 | 4.48 |
| Negative stain TEM, r (nm) | 6.01 | 6.39 | 6.84 |

**Table S2. Summary Table of separation distances calculated or measured for the three-color Cascade model.**

| | mTurq2→mVenus | mTurq2→mScarlet-I | mVenus→mScarlet-I |
|---|---|---|---|
| Energy Transfer Rates, r (nm) | 6.41 | 6.25 | 6.91 |
| AlphaFold3 model, r (nm) | 4.04 | 4.32 | 4.48 |
| Negative Stain TEM, r (nm) | 6.01 | 6.39 | 6.84 |
